# Genetically distinct neurodegenerative proteinopathies engage overlapping proteostasis components

**DOI:** 10.64898/2026.09.28.753000

**Authors:** Bhuvaneshwar Yarlagadda, Vijay Kumar

**Author notes:** **Correspondence:** Prof. Vijay Kumar, Department of Biotechnology, Anand School of Engineering and Technology, Sharda University Agra, Uttar Pradesh-282007, India.

## Abstract

Neurodegenerative proteinopathies are defined by distinct proteins whose folding, trafficking and clearance depend on shared proteostasis machinery. Whether these disorders engage common components or only common processes has been unresolved. We mapped 16 disease proteins from 12 disorders onto a fixed 990-protein proteostasis network. Across the 12 disorders, the neighborhoods of these proteins shared specific proteins beyond expectations from process composition and connectivity. The same held for the two disorders defined by more than one protein. Nine proteins contributed disproportionately, and only autophagy’s removal significantly reduced similarity. Removing PRKN, HSPA8 or UBQLN2 in silico reduced path efficiency, marking them candidate shared network vulnerabilities. In human brain proteomes, Alzheimer’s disease-zone proteins showed greater abundance remodeling in the insoluble fraction; recurrence across disorders showed no such association. The result is a component-level architecture prioritizing network-derived candidates for cross-disease functional testing.

## Introduction

Neurodegenerative diseases are an increasing global health concern, and most remain without therapies that prevent or arrest their progression.^1^ Most of these disorders are characterized, pathologically and molecularly, by disease-associated proteins and their altered assemblies: amyloid-beta^2^ and tau^3^ in Alzheimer’s disease, alpha-synuclein in Parkinson’s disease,^4^ TDP-43 and related RNA-binding proteins in amyotrophic lateral sclerosis and frontotemporal dementia (ALS/FTD),^5^ the prion protein in prion disease,^6^ and expanded-polyglutamine proteins in Huntington’s disease^7^ and the polyglutamine ataxias.^8^ Research and therapeutic development have largely followed this disease-specific, protein-focused framework. Therapies directed at individual pathological proteins have produced heterogeneous outcomes, ranging from trials that did not meet their primary efficacy endpoints to modest slowing of clinical decline in selected early-stage Alzheimer’s disease populations.^9–12^ Trial outcomes alone do not motivate the question. Although their initiating mechanisms differ, each of these disorders engages the same cellular system for protein folding, transport, quality control and clearance, and whether it engages the same components or only the same processes decides whether what is learned in one disorder can transfer to another.

Protein folding, vesicular trafficking, macroautophagy and the ubiquitin-proteasome system form an integrated proteostasis network that maintains protein localization, quality and turnover throughout the life of a post-mitotic neuron, while its functional reserve declines with age.^13–16^ This decline is a recognized hallmark of aging and a permissive condition for the age-related onset shared by these disorders. Disease-associated proteins with different structures and cellular functions are processed through this shared cellular machinery.^17^ This common dependence provides the biological basis for asking whether their network neighborhoods converge on the same individual components.

Direct experimental evidence implicates autophagy, the ubiquitin-proteasome system, chaperone activity and mitochondrial clearance across these disorders. Neuronal loss of basal autophagy causes neurodegeneration with ubiquitinated inclusions,^18–19^ autophagy induction reduces mutant huntingtin accumulation and toxicity,^20^ and expanded polyglutamine tracts impair Beclin 1-dependent autophagy.^21^ An aggregation-prone polyglutamine fragment inhibits the ubiquitin-proteasome system, evidence that disease-associated aggregation can compromise shared degradation machinery.^22^ PINK1 acts upstream of Parkin in a shared mitophagy pathway,^23–24^ mutations in the ubiquitin-processing factors VCP and UBQLN2 cause familial ALS and FTD,^25–26^ and heat-shock protein 70 suppresses alpha-synuclein toxicity in vivo.^27^ These studies indicate that proteostasis-network capacity contributes to neuronal vulnerability alongside the identities of the aggregating disease proteins. Network approaches make this engagement quantitative: disease-associated proteins occupy localized, overlapping interactome neighborhoods rather than random ones, and the degree of overlap between two diseases relates to their shared biology,^28–29^ systematic interaction screening across neurodegenerative disease proteins links them through shared partners and identifies modifiers of toxicity and aggregation,^30^ co-expression modules conserved across dementia proteomes are derived from the disease measurements themselves,^31^ comparative analyses have begun to define shared and disease-distinctive proteostasis signatures,^32^ and plasma-proteomic profiling across Alzheimer’s, Parkinson’s and frontotemporal dementia detects both shared and disease-specific alterations.^33^ The prion-like misfolding of a disease-associated protein is itself a principle shared across many of these disorders,^34^ which motivates asking whether the downstream systems handling those proteins are shared as well.

Despite this convergent evidence, it remains unresolved whether these diseases engage the same proteostasis components or only the same proteostasis processes. Pathway-enrichment analysis cannot separate the two, because two diseases can draw similar proportions of autophagy, degradation and chaperone proteins while sharing almost none of the same proteins. Similar process composition can therefore coexist with limited overlap in protein identity. Individual diseases are studied on separate, non-comparable maps, which makes it difficult to identify shared, cross-disease structure from features restricted to a single disorder, and thereby to prioritize mechanisms that might be relevant across multiple diseases. Two things follow when several diseases engage the same protein. It sits in the neighborhoods of more than one of them, so it is a point at which otherwise distinct disorders intersect. And because the diseases are genetically unrelated, its recurrence is unlikely to reflect the peculiarities of any one disorder or any one dataset. On that reasoning a shared protein becomes a candidate for comparative functional study, and establishing its contribution to shared proteostasis architecture requires comparison across diseases.

In a preceding study, we developed a disease-naive proteostasis framework and applied it to a single disease, characterizing a shared APP-MAPT convergence zone in Alzheimer’s disease.^35^ That framework organizes protein quality control into four modules defined independently of any disease data, allowing the disease-associated seed proteins of any proteinopathy to be mapped onto the same reference network. Its single-disease application established that two disease-associated seed proteins of one disease share a common proteostasis neighborhood, and it left open the comparative question addressed here, of whether genetically and clinically distinct diseases engage common components more often than their process composition and connectivity would predict.

We extended the disease-naive proteostasis framework, defined as a fixed 990-protein backbone across the same four modules, to a panel of twelve neurodegenerative diseases represented by sixteen disease-associated seed proteins. On this common backbone we constructed a proteostasis zone for each disease from the interactions of its disease-associated seed proteins, and quantified their global and pairwise organization, their differential proteostasis composition, which proteins most alter the shared structure when removed in silico, and the topological consequences of these in silico perturbations. Each inference was evaluated against a matched permutation null appropriate to the test, and the principal comparative analyses were repeated across all five STRING evidence regimes to separate structure that is robust to evidence strength from signal that depends on it. This yields a comparative proteostasis landscape on which the twelve diseases can be placed and compared (**Figure 1**). Network position, however, is not the same as molecular behavior in affected brain. We therefore additionally asked whether position within this shared architecture, meaning the set of proteostasis proteins the diseases occupy together with the connections among them, predicts molecular alteration, by analyzing a pan-neurodegeneration brain-proteome resource spanning six disorders^36^ under the same framework.

**Figure 1.**
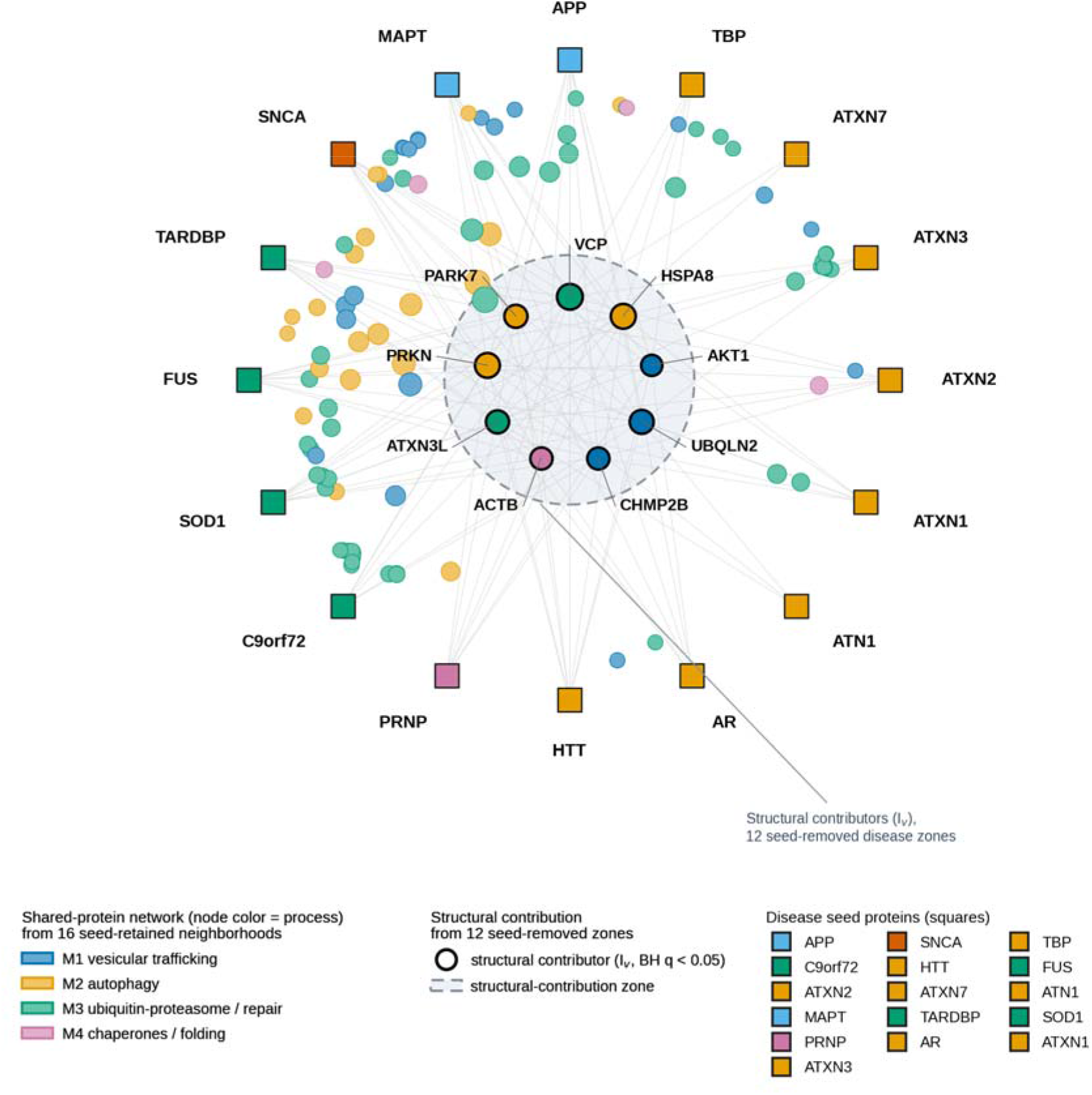
Integrative map of the sixteen disease-associated seed proteins and recurrent proteostasis proteins. Radial layout of the whole panel. The sixteen disease-associated seed proteins (squares) are arranged on the outer ring, grouped and colored by disease class (ALS/FTD proteins shown in green). Proteostasis proteins present in at least four seed-protein neighborhoods are drawn inside, colored by biological process and placed nearer the center the more disease-associated seed proteins share them. The nine proteins with a significant structural contribution (*I*_*v*_, BH *q* < 0.05) are ringed in black and labeled on the inner ring. Two distinct analytical quantities are combined here: the displayed network of shared proteins is derived from the sixteen seed-retained seed-protein neighborhoods, whereas the black structural-contribution rings are derived from the twelve primary seed-removed disease zones; structural contribution was not tested in the seed-retained universe used to draw the network. The shaded inner zone visually demarcates the seed-removed structural contributors from the surrounding seed-retained shared-protein network. See also Tables S10, S11 and S12.

## Results

### One fixed map of protein quality control for twelve proteinopathies

The fixed, disease-naive backbone comprises 990 proteins in four mutually exclusive modules: vesicular trafficking (M1, 478 proteins), autophagy (M2, 102), the ubiquitin-proteasome system and protein repair (M3, 331), and chaperones and folding (M4, 79) (Figure S1; Table S1). It was curated from Reactome pathway annotations, mapped to STRING v12.0, restricted to associations among the curated proteins, and assembled without reference to the diseases later mapped onto it,^35^ and structure detected among diseases therefore reflects where their seed proteins fall on a fixed network (Methods). Sixteen disease-associated seed proteins were mapped to twelve diseases spanning Alzheimer’s disease, Parkinson’s disease, ALS/FTD, prion disease, Huntington’s disease and seven additional polyglutamine disorders, namely spinal and bulbar muscular atrophy (SBMA), dentatorubral-pallidoluysian atrophy (DRPLA) and spinocerebellar ataxia types 1, 2, 3, 7 and 17 (SCA1, SCA2, SCA3, SCA7 and SCA17). Four of the seed proteins (APP, AR, ATXN3 and ATXN7) are themselves constituents of the proteostasis machinery and reside within the backbone; these four were removed from every eligibility-restricted analysis, leaving a primary eligible universe of 986 proteins.

We refer to each disease-associated protein used to anchor the network mapping as a seed protein, to its module-restricted set of associated backbone proteins as a seed neighborhood, and to the disease-level set built from those neighborhoods as a disease zone. For each disease we identified, module by module, the proteostasis proteins that its seed proteins associate with in STRING (Figure 2); where a disease had more than one seed protein, we kept only the proteins shared by all of them within a module, and the union of these sets across the four modules, with the seed proteins removed, formed that disease zone (Methods). A disease zone is therefore a set of proteins associated with the seed proteins of that disease in the reference network, and not a set of proteins known to be altered in that disease. Seed-removed zone sizes ranged from 5 to 136 proteins (Figure 3; Table S2). The smallest zone, DRPLA (POLYQ_ATN1, *n* = 5), is labeled low-coverage wherever it appears and is treated as descriptive only, because permutation estimates for a five-protein set are unstable. All inference used add-one permutation p-values with Benjamini-Hochberg (BH) correction as primary and Benjamini-Yekutieli (BY) as a conservative sensitivity, computed within fixed, non-pooled correction families (Methods).

**Figure 2.**
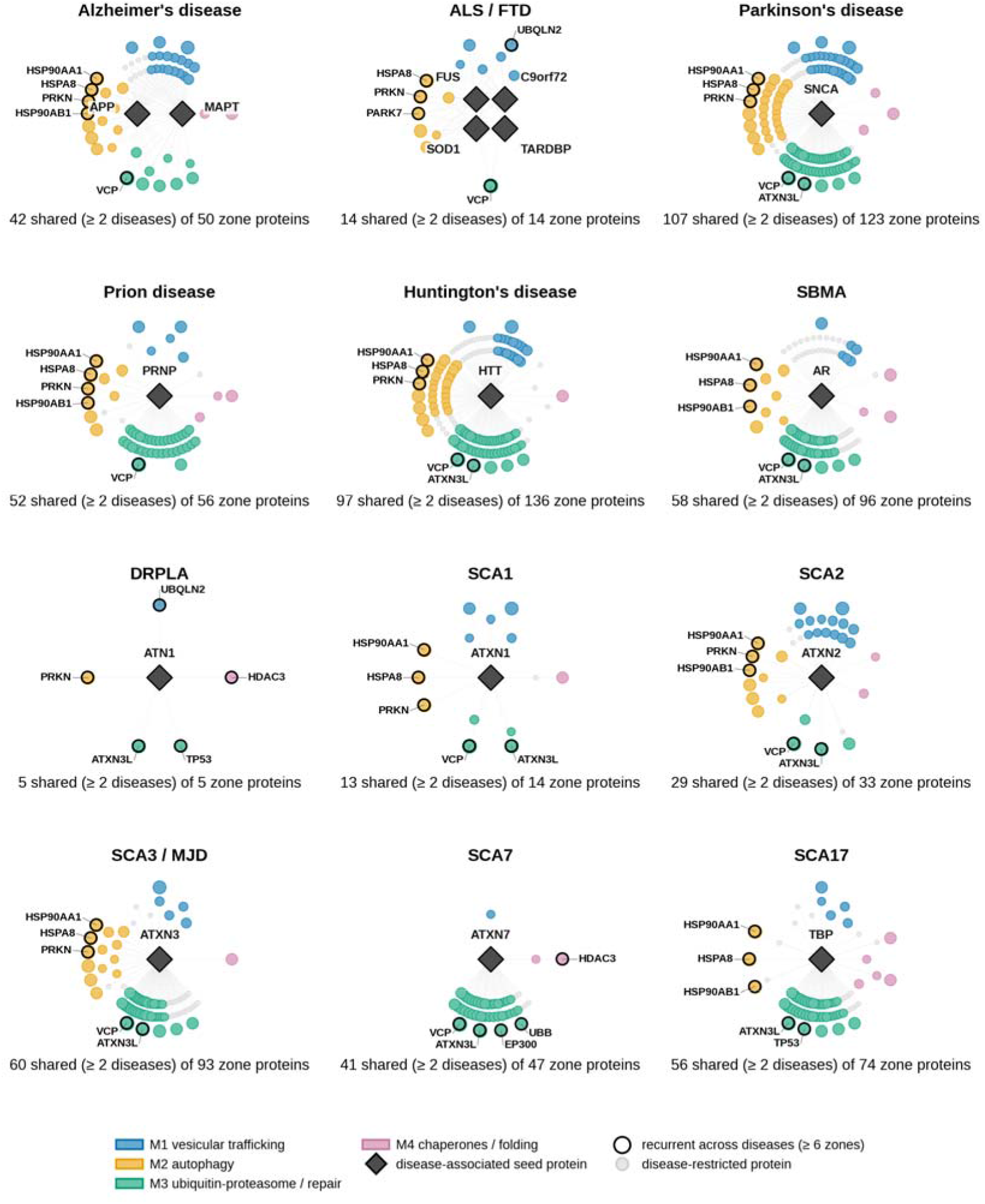
Disease-associated seed proteins engage shared and disease-restricted proteostasis neighborhoods. Twelve radial panels, one per disease. The disease-associated seed proteins (dark diamonds, labeled) sit at the center of each panel, and the proteins of that disease’s primary seed-removed zone (STRING combined score at least 0.4; eligible-universe restricted) are arranged around them by biological process: vesicular trafficking (M1, top), autophagy (M2, left), ubiquitin-proteasome processing and protein repair (M3, bottom) and chaperones and protein folding (M4, right). Marker size scales with cross-disease recurrence; proteins present in at least six disease zones are ringed and labeled as recurrent across diseases, and disease-restricted proteins (present in a single zone) are shown in gray. The count beneath each panel gives the number of zone proteins shared by at least two diseases out of the total zone size. The DRPLA zone is low coverage and is shown for descriptive purposes only.

**Figure 3.**
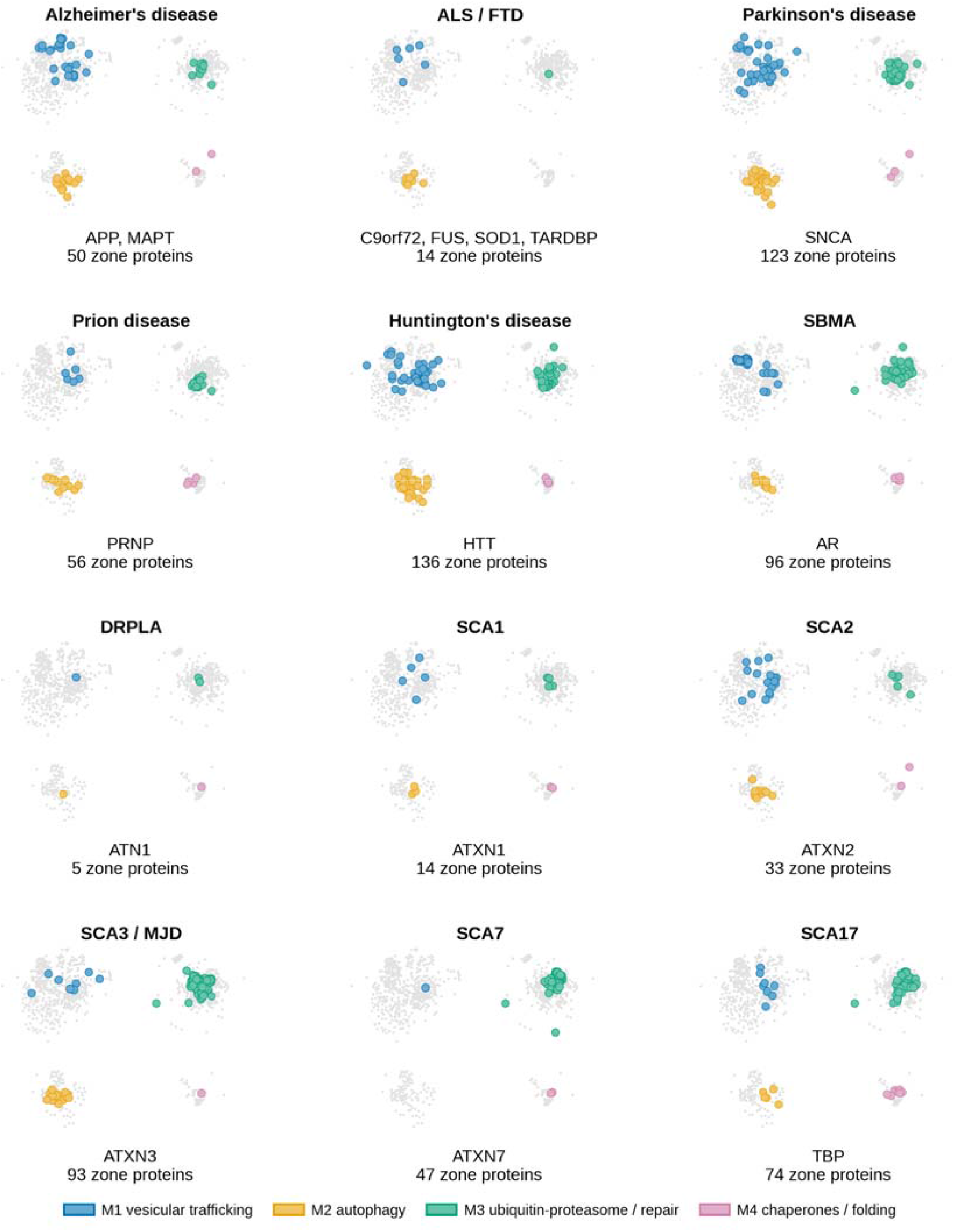
Disease-associated proteostasis neighborhoods occupy distinct regions of the shared network. Twelve panels, one per disease. The complete 990-protein backbone is shown in gray, and the proteins of each disease’s primary seed-removed zone are highlighted and colored by biological process (vesicular trafficking M1, blue; autophagy M2, orange; ubiquitin-proteasome processing and protein repair M3, green; chaperones and protein folding M4, purple). The primary evidence regime is C400 (STRING combined score at least 0.4), and only eligible-universe proteins are included; the two multi-protein diseases (Alzheimer’s disease and ALS/FTD) use the prespecified within-disease intersection rule. DRPLA is low coverage and descriptive only. See also Table S2.

### Different disease proteins of one disorder engage a common set of quality-control proteins

Ten of the twelve diseases are represented by a single seed protein, whereas Alzheimer’s disease (APP and MAPT) and ALS/FTD (TARDBP, FUS, SOD1 and C9orf72) are each represented by several. For these two we asked whether their seed proteins associate with a common set of proteostasis proteins or engage the network separately. In Alzheimer’s disease, 51 proteostasis proteins were shared between the neighborhoods of APP and MAPT, against a composition-matched null mean of 17.4 (2.9-fold; empirical *P* = 1/10,001, 10,000 permutations; Figure 4A; Table S3). In ALS/FTD, 16 proteostasis proteins were shared across the neighborhoods of all four seed proteins, against a null mean of 0.17 (empirical *P* = 1/10,001; Table S3). Because four-way intersections are intrinsically sparse, this result establishes a 16-protein common set beyond its matched expectation, and direct effect-size comparison with the two-seed Alzheimer’s disease result is inappropriate. These within-disease shared sets (51 proteins for Alzheimer’s disease, 16 for ALS/FTD) include the backbone-resident seed proteins; after the eligible-universe restriction, the primary seed-removed Alzheimer’s disease and ALS/FTD zones contain 50 and 14 proteins, respectively. The seed proteins of a multi-protein disease thus share a common proteostasis neighborhood larger than a matched null predicts, although most of what each seed protein engages is not shared: the four ALS/FTD seed neighborhoods contain 38 to 95 proteins each against a 16-protein four-way intersection. The Alzheimer’s disease and ALS/FTD zones used in the comparisons below represent this shared neighborhood. Proteins as different as APP and tau, or TDP-43 and SOD1, are therefore associated with an overlapping set of quality-control proteins rather than with separate ones.

**Figure 4.**
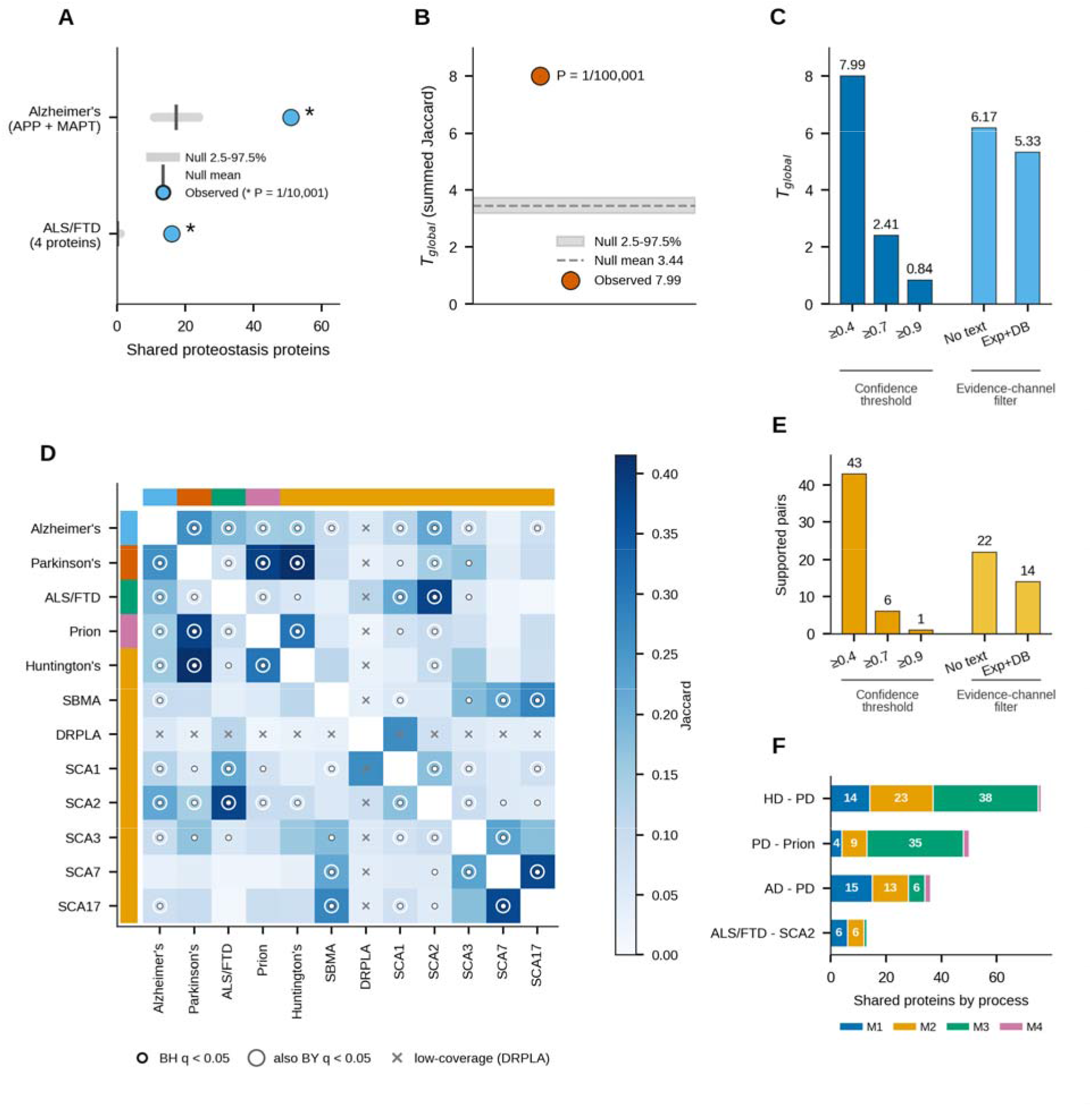
Neurodegenerative diseases form a non-random proteostasis structure. (A) Within-disease overlap: proteostasis proteins shared across the multiple disease-associated seed proteins of Alzheimer’s disease (APP, MAPT) and ALS/FTD (four disease-associated seed proteins); the observed count is shown against the null mean and the null 2.5th-97.5th-percentile interval (^*^empirical *P* = 1/10,001). (B) Observed global disease-zone similarity (summed pairwise Jaccard, *T*_global_) at C400 against the matched null (*P* = 1/100,001). (C) The global similarity statistic across the five STRING evidence regimes, shown as bars grouped into ordered confidence thresholds (C400, C700, C900) and alternative evidence-channel filters (NONTEXT, EXPDB) rather than as a single continuous series. (D) Pairwise disease-zone Jaccard similarity at C400 with statistical-support symbols (filled dot, BH q < 0.05; filled dot within an open ring, also BY q < 0.05; cross, low-coverage DRPLA pair); side bars indicate disease class. (E) Number of formally supported pairs across the same five regimes, grouped as in (C). (F) Biological-process composition of the shared proteins of selected cross-class relationships, by process. See also Table S4 and Figure S3.

### Twelve proteinopathies engage overlapping sets of individual proteostasis proteins

The twelve diseases engaged overlapping sets of individual proteins far more often than zones matched for process composition and connectivity would predict. We summarized the panel with a single global statistic, the summed pairwise Jaccard similarity across all 66 disease pairs, tested against a joint null that independently randomizes each zone while preserving its size and module-by-degree-bin composition (100,000 permutations; Methods). In the primary evidence regime (STRING combined score at least 0.4; C400) the observed global similarity greatly exceeded the null (*T*_global_ = 7.99, against a null mean of 3.44 with a 2.5th-to-97.5th-percentile interval of 3.19 to 3.73, from 100,000 permutations; empirical *P* = 1/100,001) (Figure 4B; Table S4). This exceedance was not attributable to any single disease or to the largest zones, and excluding DRPLA retained non-random global overlap in a 10,000-permutation sensitivity (*T*_global_ = 7.22 against a null mean of 3.28; empirical *P* = 1/10,001; Figure S2A, S2B; Table S4). STRING aggregates evidence channels of differing directness, from co-expression and text-mining to curated database and direct experimental support, and we recomputed the global statistic under five prespecified regimes that progressively restrict the admissible evidence. The NONTEXT and EXPDB regimes filter associations on the evidence channels they carry and do not recompute the combined score with text mining removed, so they test whether the result rests on text co-mention alone rather than removing literature bias from the network. The global statistic remained above its null in every regime, with *T*_global_ = 2.41 against a null mean of 0.94 under C700 (score at least 0.7), 0.84 against 0.25 under C900 (score at least 0.9), 6.17 against 2.90 under a regime requiring non-text-mining support (NONTEXT), and 5.33 against 2.57 under an experimental- or database-supported regime (EXPDB); the empirical *P* was 1/100,001 in every regime except C900, where it was 6/100,001 (Figure 4C; Table S4). The absolute statistic falls as the zones contract. Along the confidence-threshold axis the ratio of observed to expected similarity rises, from 2.3 in the primary regime to 2.6 under C700 and 3.3 under C900, whereas under the two evidence-channel regimes it is slightly lower than in the primary regime, 2.1 under NONTEXT and 2.1 under EXPDB. Disease-zone sizes contracted as thresholds tightened (Figure S3A; Table S2), and the NONTEXT and EXPDB results show that the global conclusion does not rest on associations supported by literature co-mention alone.

Because eight of the twelve diseases are polyglutamine disorders, accounting for 28 of the 66 disease pairs, we tested directly whether this global overlap was driven by this single mechanistic family; and it was not. Restricting the statistic to the 38 pairs that are not internal to the polyglutamine family retained strong overlap (*T*_cross_ = 4.75 against a null mean of 1.93 with a 2.5th-to-97.5th-percentile range of 1.74 to 2.14; empirical *P* = 1/100,001), and these cross-class pairs accounted for the majority of the global statistic (59 percent, versus 41 percent from within-polyglutamine pairs). Leaving out any single disease left the panel strongly non-random (all twelve leave-one-disease-out statistics *P* = 1/100,001 at 100,000 permutations), as did every family-balanced five-disease panel formed from the four non-polyglutamine diseases and a single polyglutamine representative (all eight representatives *P* = 1/100,001) (Table S5). The comparative overlap is therefore a property of the panel as a whole and not an artifact of polyglutamine-family over-representation. The excess overlap is therefore not explained by a general reliance on protein quality control, which would be expected of any aggregating protein, but by recurrence of the same individual proteins within it.

### Sharing crosses amyloid, synuclein, prion and polyglutamine disease classes

Most of the panel participates in this sharing. Of the 66 disease pairs, 43 were formally supported in the primary regime, meaning the global gate was passed and the pairwise BH q was below 0.05. Setting aside the seven supported DRPLA comparisons leaves 36 of 55 pairs, 23 of them joining diseases from different genetic or clinical classes (Figure 4D, 4E; Table S4). The sharing extends well beyond disorders that resemble each other mechanistically. Huntington’s disease and Parkinson’s disease shared 76 proteins (Jaccard 0.415; null mean 0.186); Parkinson’s disease and prion disease shared 50 proteins (Jaccard 0.388; null mean 0.148); and Alzheimer’s disease and Parkinson’s disease shared 36 proteins (Jaccard 0.263; null mean 0.058); the biological-process composition of the shared proteins in these pairs is shown in Figure 4F (Table S4). A strongly supported cross-class relationship with established biological context joined ALS/FTD and SCA2 (POLYQ_ATXN2), which shared 13 proteins (Jaccard 0.382; BH *q* = 4.7×10^−5^) with equal representation of vesicular trafficking (M1) and autophagy-centered quality control (M2), six proteins in each, and a single ubiquitin-processing protein (M3) (Figure 4F); the observed-versus-null comparison for this and the other prespecified selected pairs is shown in Figure S3B.

The strength of individual pairwise relationships depended on evidence stringency. The number of formally supported pairs fell from 43 in the primary regime to 22 under NONTEXT, 14 under EXPDB, 6 under the higher-confidence C700 threshold and 1 under the highest-confidence C900 threshold (Figure 4E; Table S4). This attenuation is consistent with contraction of disease zones under stricter evidence criteria, because tighter thresholds shrink each zone and consequently reduce the overlap available for any pair. Every pair supported under a restricted regime was also supported in the primary regime, and the supported sets are strictly nested, the C900 set within the C700 set, the EXPDB set within the NONTEXT set, and both within the primary set (Table S4). Four relationships were supported in at least three of the four restricted regimes, joining Alzheimer’s disease with Parkinson’s disease, Huntington’s disease with Parkinson’s disease, SBMA with SCA17, and SCA7 with SCA17, the last being the only pair supported in all five regimes (Table S4). These four are the most evidence-robust relationships in the panel. The global architecture holds under every regime, whereas any one pair outside those four is evidence-weighted and should be read as hypothesis-generating. A descriptive dendrogram summarizes the primary-regime similarity structure (Figure S2C).

### Disease neighborhoods draw disproportionately on autophagy and ubiquitin-dependent degradation

Most diseases weight the four processes unevenly rather than drawing from them in proportion. Testing this requires a null that preserves the degree distribution of each zone but not its proteostasis-process membership, against which the composition of each zone is compared module by module and summed into one statistic per disease (Methods). In the primary regime, 10 of 12 diseases showed a significant omnibus deviation in relative proteostasis composition (Figure 5A; Table S6); the count was 8, 7, 9 and 9 under the C700, C900, NONTEXT and EXPDB regimes, respectively; this compositional organization, like the global signal, is broadly robust to evidence restriction (Figure S4A, S4B). At the process level, autophagy (M2) was positively represented in seven of twelve disease zones (Alzheimer’s disease, ALS/FTD, Huntington’s disease, Parkinson’s disease, SCA2, SCA3 and prion disease), and the ubiquitin-proteasome and protein-repair module (M3) was positively represented in five of twelve disease zones (SBMA, SCA3, SCA7, SCA17 and prion disease) (Figures 5B and S2D; Table S7). The principal autophagy and ubiquitin-processing patterns were retained in the NONTEXT and EXPDB networks; stricter confidence thresholds reduced testability and altered several individual disease-process contrasts, indicating that process-level inference is more sensitive to network sparsity than the global overlap result (Table S7). Because the four module proportions of a zone are interdependent, over-representation in one process necessarily lowers the relative representation of others; conditioned on module sizes and degree, the negative direction for vesicular trafficking (M1) is a compositional consequence of enrichment in the degradation-associated processes and is not interpreted as loss or impairment of vesicular trafficking. Diseases show proportionally greater representation in the degradation-associated processes along two overlapping axes defined by proteostasis process, an autophagy-weighted axis and a ubiquitin-processing-weighted axis. The axes overlap, and two diseases, SCA3 and prion disease, are represented on both (Figure 5C). Autophagy and ubiquitin-dependent degradation are the two routes by which a cell disposes of material it cannot refold, and it is these two that the disease neighborhoods draw on most.

**Figure 5.**
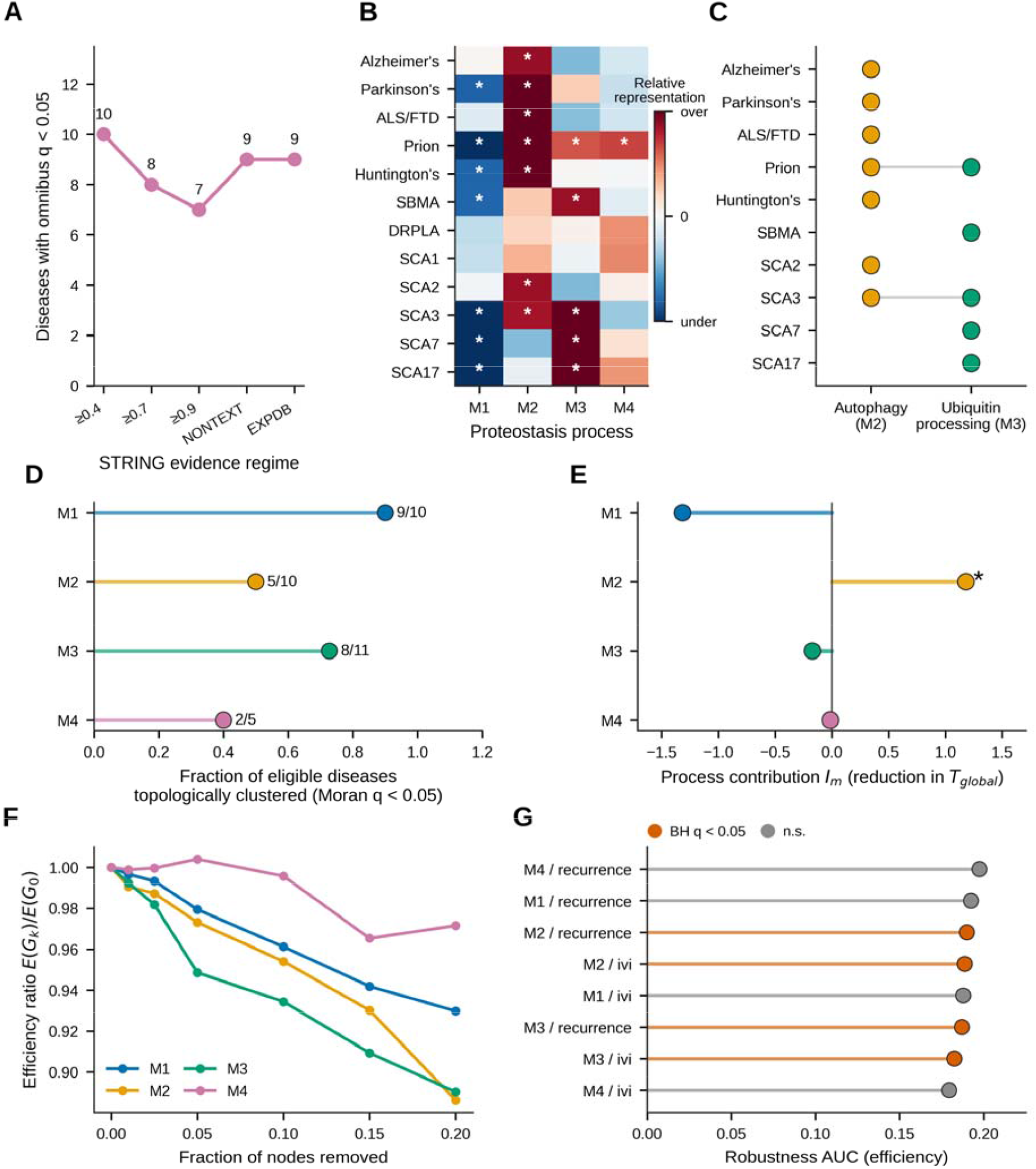
Proteostasis processes differ in their disease composition, topological clustering, contribution to shared structure and robustness to node removal. (A) Number of diseases with a significant omnibus compositional deviation across regimes. (B) Relative process representation per disease at C400, shown as a signed deviation from the composition-matched expectation (red over-represented, blue under-represented; color intensity scaled by the magnitude of the deviation; asterisks, BH *q* < 0.05). (C) The two overlapping process axes: diseases positively represented in autophagy (M2) and in ubiquitin-proteasome processing and protein repair (M3). (D) Fraction of testable diseases whose zone proteins are topologically clustered within each process-specific network (Moran statistic, BH *q* < 0.05; counts shown as clustered over testable). (E) Process (module) contribution *I*_*m*_ to global disease-zone similarity; asterisk, BH *q* < 0.05. (F) Network robustness, the efficiency ratio under recurrence-ranked node removal for each process. (G) Robustness area-under-curve tests for efficiency across processes and rankings; vermilion, BH *q* < 0.05. **See also Figure S4 and Tables S6-S8, S13 and S15**.

### Shared trafficking involvement is concentrated on routes toward degradative compartments

The proteins a disease engages within a module sit together in particular neighborhoods rather than scattered across it. For each disease and module, we measured the topological autocorrelation of zone membership over the module interaction graph (the Moran statistic, *I*), against a membership-preserving permutation null (Methods). Membership was positively autocorrelated in 24 of 36 testable disease-module combinations after correction (BH *q* < 0.05), across all four modules (Figure 5D; Table S8), with the disease-by-module detail in Figure S2E. Clustering was most pronounced in the ubiquitin-proteasome and protein-repair module, whose mean Moran statistic across testable diseases (0.132) is more than twice that of any other module; autophagy has the lowest mean (0.046) and the fewest significant combinations of the three testable-rich modules (Table S8). The Huntington’s disease zone was clustered within the ubiquitin-proteasome and protein-repair module (M3; *I* = 0.145; BH *q* = 0.001) and within vesicular trafficking (M1; *I* = 0.088; BH *q* = 0.001), and the Alzheimer’s disease zone within vesicular trafficking (M1; *I* = 0.070; BH *q* = 0.001) and autophagy (M2; *I* = 0.074; BH *q* = 0.014).

The strongest clustering and the most consistent clustering are not in the same module. Clustering was most consistent in vesicular trafficking, the module whose relative representation was negative in the composition analysis, where 9 of the 10 testable disease combinations were significant. A disease can therefore take a relatively small share of its zone from trafficking and still concentrate that share within particular neighborhoods of the module, which makes the negative compositional direction for M1 a description of selective involvement of parts of the trafficking machinery rather than its exclusion.

A complementary prespecified test of whether these zones are enriched for the most topologically central proteins (top-decile within-module influence, measured as the Integrated Value of Influence, IVI) found no significant enrichment in any of the 36 eligible disease-module combinations after correction (minimum BH *q* = 0.33; 1,000 permutations per combination; Table S9), and a secondary test on mean and median influence was likewise non-significant (Table S9). Four combinations reached an uncorrected P below 0.05 with fold enrichment between 1.4 and 2.4, so the test bounds rather than excludes a preference for the most influential proteins at this permutation count. The shared signal therefore does not require the most influential proteins of a module, and the identity of the shared trafficking proteins indicates which part of the module is involved. The trafficking proteins recurring in four or more disease zones are CHMP2B, UBQLN2, AKT1, DCTN1, EGFR, SH3GL2 and SYNJ1, with BIN1, DNM1, DYNC1H1, KIF5A, KLC1 and KLC2 among those in three (Table S10). These belong to endocytosis and endolysosomal sorting and to microtubule-based transport rather than to bulk secretory traffic, which places the shared trafficking involvement at the steps that carry cargo toward degradative compartments. These are the steps a neuron uses to move cargo from its processes to the degradative compartments of the cell body.

### Autophagy and a small set of degradation and chaperone proteins carry the shared overlap

Eighteen proteins occur in at least six of the twelve zones (Table S10). The most recurrent proteins with a well-characterized proteostasis-effector role were VCP (present in 10 of 12 diseases); HSPA8, PRKN, HSP90AA1 and ATXN3L (9 of 12); HSP90AB1 and UBQLN2 (8 of 12); PARK7 and PINK1 (7 of 12); and CHMP2B and SQSTM1 (6 of 12) (Figure 6A, 6B; Table S10). A complementary count across the sixteen individual seed-protein neighborhoods, with the seed proteins retained, gave a concordant ranking, led by VCP and ataxin-3 (ATXN3), each in fourteen of sixteen, and PRKN, HSPA8 and HSP90AA1, each in thirteen (Figure 6C; Table S11). Here ATXN3L denotes ataxin-3-like, a paralog of the SCA3 disease protein ataxin-3 (ATXN3); ATXN3L is a constituent of the eligible proteostasis universe and is distinct from ATXN3, which is a seed protein removed from the seed-removed zones. These proteins act at successive points on the same substrate path. VCP extracts ubiquitinated proteins from membranes and complexes so that they can be handed on, and UBQLN2 shuttles ubiquitinated substrates to the proteasome. PRKN and PINK1 act together in mitochondrial quality control, alongside PARK7. SQSTM1 binds ubiquitinated cargo and links it to the forming autophagosome, and CHMP2B, a subunit of the endosomal sorting complex required for transport (ESCRT-III), closes and matures the endosomal and autophagic membranes that carry it. The chaperones HSPA8, HSP90AA1 and HSP90AB1 hold misfolded clients in a soluble state. Between them these proteins cover recognition, extraction, delivery and folding, the successive steps by which a cell disposes of a protein it cannot repair. Recurrence counts how often a protein appears; it does not establish that its presence contributes to cross-disease similarity, and we quantified the structural contribution of each eligible protein as *I*_*v*_, the reduction in global similarity when that protein is removed from every zone, evaluated against the same C400 permutation null (Methods). That null already expects a widely shared protein to remove more similarity when it is deleted, so a significant Iv marks a protein that exceeds the expectation set by its own recurrence rather than one that is simply recurrent.

**Figure 6.**
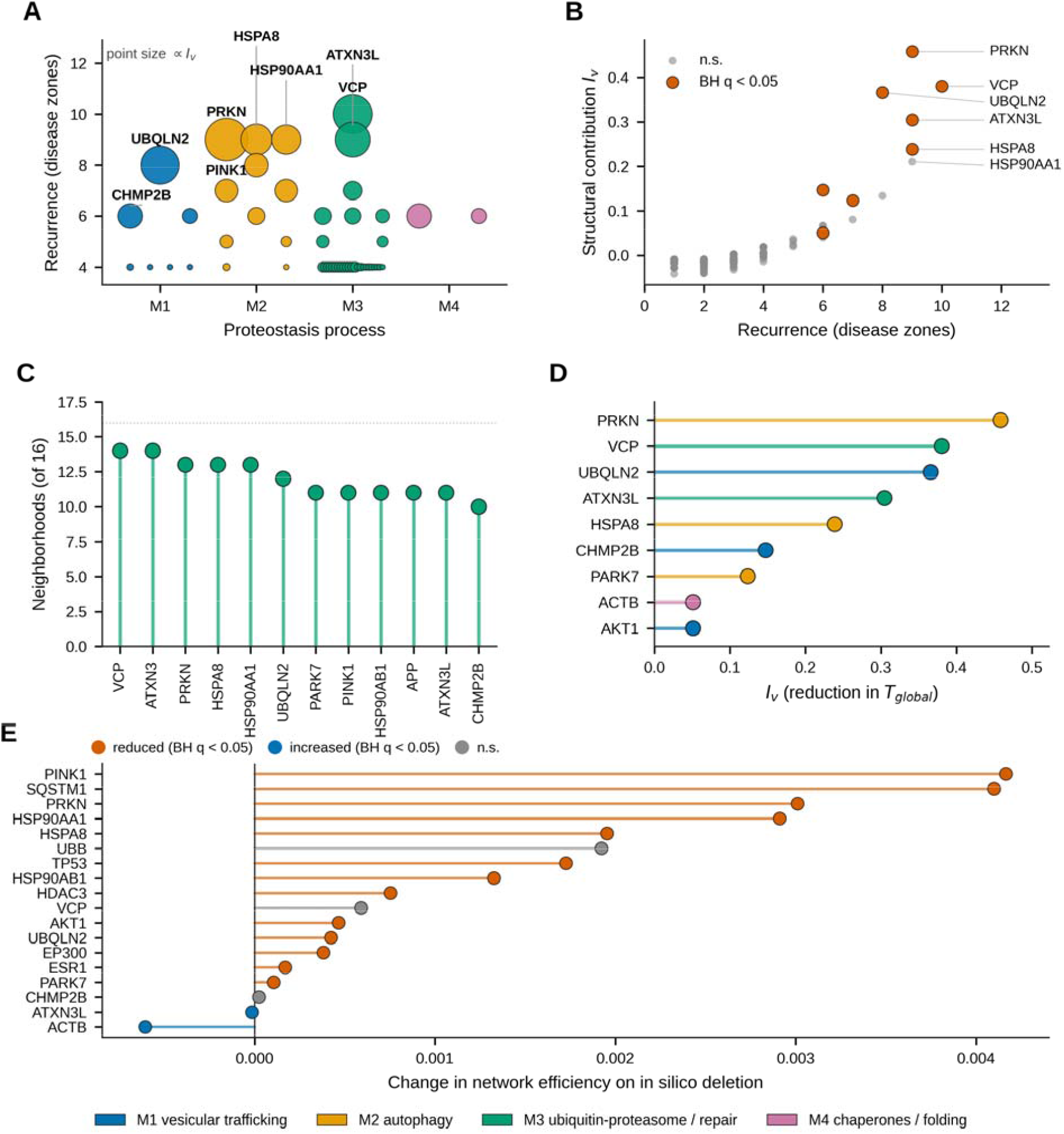
Recurrent proteostasis proteins and their structural contribution. (A) Recurrent proteins (present in at least four disease zones) arranged by proteostasis process (M1 vesicular trafficking, M2 autophagy, M3 ubiquitin-proteasome and repair, M4 chaperones); point size scales with structural contribution *I*_*v*_. (B) Recurrence (number of disease zones) versus structural contribution *I*_*v*_ for eligible proteins; proteins with significant *I*_*v*_ (BH *q* < 0.05) are shown in vermilion and leading proteins are labeled. (C) Cross-seed recurrence, the number of seed-protein neighborhoods (of 16) in which each protein appears. (D) The nine proteins with significant structural contribution *I*_*v*_, colored by process. (E) Change in network efficiency on single-node in silico ablation for the eighteen cross-disease candidates (vermilion, significant reduction; blue, significant increase; gray, not significant). See also Figure S3.

UBQLN2, CHMP2B, VCP, ATXN3L, PARK7, ACTB, AKT1, HSPA8 and PRKN each contributed individually to cross-disease overlap, nine of the 986 eligible proteins whose contributions remained significant after correction (Figure 6B, 6D; Table S12). The largest structural contributions were PRKN (*I*_*v*_ = 0.458), VCP (0.381), UBQLN2 (0.366), ATXN3L (0.305) and HSPA8 (0.239). Seven of the nine remain significant under the conservative BY correction; PRKN (BY *q* = 0.074) and HSPA8 (BY *q* = 0.055) do not. ATXN3L is the least broadly distributed of the nine, its nine zones spanning two disease classes against four or five for the others, so its contribution is largely internal to the polyglutamine family. Recurrence and structural contribution separate more sharply than a shared ranking would suggest. Several of the most recurrent proteins did not exceed that expectation: HSP90AA1 in nine zones (*I*_*v*_ = 0.211; BH *q* = 0.62), PINK1 in seven (*I*_*v*_ = 0.126; *q* = 1.00), TP53 in seven (*I*_*v*_ = 0.081; *q* = 0.93) and SQSTM1 in six (*I*_*v*_ = 0.067; *q* = 0.21). Three proteins present in only six zones were among the most significant contributors: CHMP2B (*I*_*v*_ = 0.147; *q* = 0.0025, with no permutation of 100,000 reaching the observed value), ACTB and AKT1 (*I*_*v*_ = 0.051 for both; *q* = 0.0033 and 0.0042), the last two occupying identical zone sets and therefore statistically indistinguishable. The magnitude of *I*_*v*_ tracks recurrence closely (Spearman *ρ* = 0.84 across the 174 proteins present in at least two zones), and the three six-zone contributors have the smallest values of the nine. The two quantities separate on significance rather than magnitude, because the null scales its expectation with recurrence, so ranking by recurrence alone would not have selected this set. Three of the nine significant contributors, UBQLN2, CHMP2B and AKT1, belong to the vesicular trafficking module whose relative representation is negative, consistent with the clustering result above: the trafficking involvement of these diseases is narrow but structurally weighted. Removal of autophagy (M2) significantly reduced global cross-disease similarity in the primary analysis (*I*_*m*_ = 1.18); removal of the other modules did not pass correction (Figure 5E; Table S13). Autophagy is therefore the process on which the shared structure most depends, and the proteins that carry it are those a cell uses to recognize, deliver and degrade material it cannot refold. The seed proteins that share these proteins, together with the nine significant structural contributors among them, are summarized in an integrative protein-sharing map (Figure 1); the disease pairs sustained by each leading protein are shown in Figure S3C.

### Single-node ablation identifies proteins associated with reduced module efficiency

Removing a single shared protein from its module graph reduced the efficiency with which paths connect the remaining proteins. Of 18 cross-disease candidates tested, 15 showed efficiency changes exceeding their rewired-network null distributions after BH correction, thirteen of them positive, indicating reduced efficiency after node removal, and two small and negative (Methods). The thirteen positive changes span 1.1×10^−4^ to 4.2×10^−3^ in efficiency and comprise PINK1, SQSTM1, PRKN, HSP90AA1, HSPA8, TP53, HSP90AB1, HDAC3, AKT1, UBQLN2, EP300, ESR1 and PARK7 in descending order of effect. Removing a single candidate left the largest-component fraction unchanged in every case, the observed change being exactly zero for all 18, because no candidate is an articulation point of its module graph (Figure 6E; Table S14). Eight of the 15 efficiency effects remain significant under the conservative BY correction. No single one of these proteins holds its module together; each instead lies on many of the short routes through it. The connectivity statistic could not move for any single node tested here, so the efficiency result stands on its own.

At the module level, targeted sequential removal produced a related pattern. Sequential removal of proteins ranked by recurrence or by within-module influence degraded the efficiency of the autophagy and ubiquitin-processing modules faster than matched-random removal. The robustness area-under-curve differences were significant for autophagy (recurrence-ranked efficiency BH *q* = 0.0008; influence-ranked efficiency BH *q* = 0.036) and for ubiquitin processing (recurrence-ranked efficiency BH *q* = 0.015; influence-ranked efficiency BH *q* = 0.0008) (Figure 5F, 5G; Figure S4C, S4D; Table S15). Effects on the largest-component fraction were not significant, with the exception of an influence-ranked ubiquitin-processing difference that was statistically detectable though extremely small, and which we do not interpret as biologically meaningful fragmentation.

### Detected divergence is explained more consistently by process weighting than by disease-restricted proteins

Only one disease showed even a weak excess of proteins confined to its own zone. Using single-zone proteins, meaning proteins that appear in only one disease zone, SCA3 (ATXN3) was the only disorder with such an excess once zone size, proteostasis composition and degree were controlled (33 observed against a null mean of 23.47; BH *q* = 0.037), and a descriptive 28 of the 33 fall in the ubiquitin-proteasome and protein-repair module (M3), consistent with the deubiquitinating function of ATXN3.^37^ The excess is threshold-dependent and does not survive conservative correction or the restricted evidence regimes (Figure S2F; Figures S3D, S3E and S3F; Table S16). The ATXN3-associated ubiquitin-processing neighborhood is therefore hypothesis-generating, and the divergence between these diseases lies in how they weight shared processes rather than in sets of disease-unique proteins.

### Alzheimer’s disease-zone proteins show greater abundance remodeling in the insoluble brain proteome

Network position and molecular behavior in affected brain are separate measurements. We analyzed the Pan-Neurodegeneration Atlas (PanNDA), a resource of post-mortem brain proteomes from six neurodegenerative disorders^36^, under the same disease-naive framework, asking whether the shared architecture is accompanied by molecular alteration (Methods). Three of the twelve diseases have both a disease zone here and a matched proteome in PanNDA (Alzheimer’s disease, Parkinson’s disease, and FTLD-TDP, frontotemporal lobar degeneration with TDP-43 pathology, aligned to ALS/FTD), and 793 of the 986 backbone proteins were quantified in at least one whole proteome (Figure 7A). We first tested whether disease-zone proteostasis proteins are preferentially altered in the matched brain. In Alzheimer’s disease, zone membership was associated with greater whole-proteome abundance remodeling than the rest of the measured backbone (*β* = 0.38 in rank-normalized units; 100,000-permutation *P* = 0.011, BH *q* = 0.03; Figure 7B-7D; Table S17), and an association was also observed in the insoluble protein fraction of the Alzheimer’s brain (*β* = 0.42; 100,000-permutation *P* = 0.002, *q* = 0.007). The corresponding associations in FTLD-TDP and Parkinson’s disease did not pass correction in either layer; in FTLD-TDP the whole-proteome estimate was *β* = 0.47 (*q* = 0.09) and the insoluble estimate *β* = 0.13 (*q* = 0.76).

**Figure 7.**
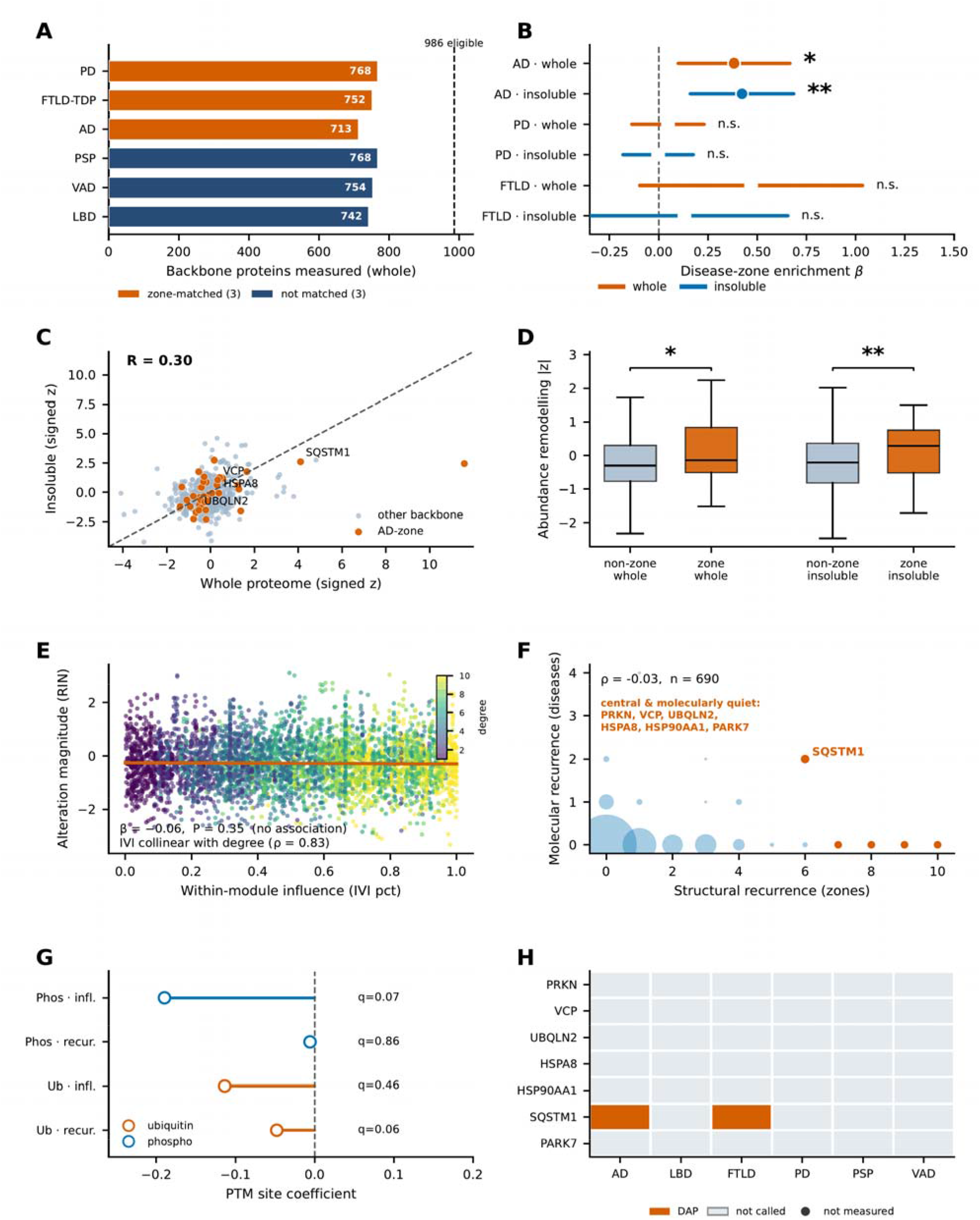
Brain-proteomic evaluation of network-defined disease zones (integrated PanNDA analysis). (A) Backbone coverage across the six PanNDA brain proteomes; the three zone-matched disorders (Alzheimer’s disease, Parkinson’s disease and FTLD-TDP) are shown in orange and the three unmatched disorders in navy, against the 986 eligible backbone proteins. (B) Disease-zone engagement: enrichment of disease-zone proteins for abundance remodeling in the matched brain, whole and insoluble layers; *β* with 95% confidence interval; filled markers and asterisks denote significance (^*^*q* < 0.05, ^**^*q* < 0.01). (C) Whole versus insoluble signed alteration for Alzheimer’s disease backbone proteins; Alzheimer’s disease-zone proteins in orange, Pearson *R* shown. (D) Abundance remodeling in Alzheimer’s disease-zone versus non-zone proteins, whole and insoluble; permutation P, with asterisks denoting ^*^*P* < 0.05 and ^**^*P* < 0.01. (E) Within-module influence (IVI percentile) versus alteration magnitude across the six proteomes; the association is flat (*β* = −0.06, *P* = 0.35) and points are colored by within-module degree, which is collinear with IVI (*ρ* = 0.83). (F) Structural recurrence versus molecular recurrence (*ρ* = −0.03, *n* = 690); selected recurrent proteins labeled. (G) Post-translational-modification site remodeling versus recurrence and influence; none pass correction. (H) Differential-abundance status of the selected recurrent proteins across the six proteomes. See also Figure S5.

Because this estimate rests on 44 zone proteins in the whole proteome and 41 in the insoluble fraction, we refit each model under two removals (Table S17). The first removed any paralogue family contributing three or more members to a zone; in the Alzheimer’s disease zone this removed eight alpha-tubulins, present because the zone is the module-restricted neighborhood of APP and MAPT. The whole-proteome estimate fell to *β* = 0.23 with permutation *P* = 0.13, while the insoluble-fraction estimate held at *β* = 0.36 with *P* = 0.014. The same rule removed 40 proteasome, RAB and kinesin-1 members from the Parkinson’s disease zone, moving its whole-proteome estimate from *β* = 0.05 to *β* = 0.22 (*P* = 0.06), and removed nothing from the ALS/FTD zone. The second removal took zone proteins one at a time, and then the three whose removal moved the estimate furthest toward zero: no single removal raised the model *P* above 0.016 in either Alzheimer’s layer, and removing GFAP, SQSTM1 and SYT11 together reduced the whole-proteome estimate to *β* = 0.24 with permutation *P* = 0.13, while the insoluble-fraction estimate fell to *β* = 0.32 with *P* = 0.02 on removing SYT1, SQSTM1 and GFAP.

The whole-proteome association therefore does not survive removal of one paralogue family, while the insoluble-fraction association does. As a disease control we refitted the same models on the PsychENCODE schizophrenia and bipolar-disorder whole proteomes, measured on the same platform and designated disease controls by the source study. No zone was associated with alteration magnitude in either disorder, at a coverage comparable to the Alzheimer’s comparison (771 backbone proteins, 44 in the Alzheimer’s disease zone): the Alzheimer’s disease zone gave *β* = 0.13 in schizophrenia and *β* = 0.02 in bipolar disorder, against *β* = 0.38 in Alzheimer’s brain, with permutation *P* = 0.39 and 0.94 (Table S17). These cohorts carry no insoluble layer, so the insoluble-fraction association could not be tested this way.

How much a protein changed in the diseased brain did not track its position in the shared architecture. Across the six disease proteomes, neither within-module influence nor disease-zone recurrence was detectably associated with alteration magnitude (influence *β* = −0.06, cluster-robust *P* = 0.35; permutation *P* = 0.36; Figure 7E; Table S17). Network influence is collinear with within-module degree, so its estimate is not separable from connectivity and is not interpreted directionally (Methods). Structural recurrence and molecular recurrence were not detectably associated (Spearman *ρ* = −0.03, *P* = 0.47; *n* = 690; Figure 7F), and within-module influence was not detectably associated with whether a protein was called altered in any disease (Fisher’s exact *P* = 0.56). Neither phosphorylation nor ubiquitination-site remodeling was associated with structural recurrence after correction; the ubiquitination-recurrence estimate was negative and did not pass the defined multiple-testing threshold (parent-protein-level permutation *P* = 0.03, BH *q* = 0.06, and *q* = 0.06 under an independent second-seed replication; Figure 7G; Table S17). The corresponding whole-versus-insoluble comparisons, architecture-state coefficients and layer interaction are shown in Figure S5A-S5D.

The selected recurrent proteins also differed in their molecular responses. UBQLN2 and VCP were not called differentially abundant in any of the six proteomes, SQSTM1/p62 was called in Alzheimer’s disease and FTLD-TDP, and the chaperones HSPA8 and HSP90AA1 were not called differentially abundant under the gene-level canonical-accession mapping, although one HSPA8 proteoform met the criterion in Parkinson’s disease, a heterogeneous rather than shared molecular response (Figure 7H). A protein can therefore be central to how these diseases are organized on the network without being among those whose abundance changes most in end-stage tissue. The published pathology-correlation files contain filtered lists of significant correlates rather than complete tested backgrounds; we therefore mapped these proteins descriptively onto the proteostasis architecture (several high-influence backbone proteins are among the TDP-43-insoluble correlates) and did not perform enrichment tests or treat proteins absent from the lists as noncorrelates (Table S18). Data sources, provenance and differential-abundance extraction quality control for this arm are reported in Tables S19 and S20, and Monte Carlo precision, independent replications and the complete statistical specification of every test in Tables S21 and S22. The complete primary C400 disease-zone networks for all twelve diseases are provided in Figures S6-S17.

## Discussion

We mapped sixteen disease-associated proteins from twelve disorders onto a network fixed before any disease was placed on it, and tested their overlap against a matched permutation null. We show that genetically distinct proteinopathies shared individual proteostasis components more extensively than expected from their process composition and network connectivity. Autophagy and selected recurrent proteins contributed to the similarity of these disease-associated neighborhoods. Candidates for functional investigation were identified on two counts: a significant reduction in cross-disease similarity when the protein was removed from every zone, and a significant reduction in path efficiency when it was removed from its module graph. The null holds the module composition and degree distribution of each zone fixed, and the excess therefore lies in the identity of the proteins occupied and not in the size, process composition or centrality of the regions occupied. The same component-level organization was present within the two disorders represented by multiple disease-associated proteins. Within both Alzheimer’s disease and the ALS/FTD spectrum, disease-associated proteins with distinct normal functions and genetic causes converged on common proteostasis neighborhoods beyond matched expectations: APP and MAPT in Alzheimer’s disease, and TARDBP, FUS, SOD1 and C9orf72 in ALS/FTD. Our preceding study characterized APP-MAPT convergence in Alzheimer’s disease.^35^ The present analysis identifies a second within-disease convergence in ALS/FTD before extending the comparison across diagnoses.

This inference is limited to Alzheimer’s disease and ALS/FTD because the other ten disorders are represented by one disease-associated protein each.

The diseases do not draw on the four processes evenly, and the pattern of that unevenness follows established biology. Autophagy was positively represented in most disease zones and was the only process whose removal reduced global similarity. Neuronal autophagy loss causes neurodegeneration with ubiquitinated inclusions,^18–19^ and autophagy induction or impairment modulates polyglutamine toxicity,^20–21^ which places autophagy at the center of the shared architecture. The ubiquitin-proteasome and protein-repair process forms a second axis, positively represented in prion disease and in four of the eight polyglutamine disorders (SBMA, SCA3, SCA7 and SCA17), with SCA1 and SCA2 negative in direction though not significant, and carrying several of the strongest protein-level structural contributions, consistent with the direct inhibition of proteasome function by aggregation-prone polyglutamine proteins.^22^ Several disorders, including SCA3 and prion disease, are represented on both axes, which makes the two overlapping dimensions of one architecture and not alternative classes. Vesicular trafficking contributed through a selective spatial organization. Although it constituted a smaller fraction of disease zones than expected, the involved proteins clustered repeatedly within endocytic, endolysosomal-sorting and microtubule-transport neighborhoods, concentrating cross-disease convergence along routes that deliver cargo toward degradative compartments. The shared trafficking signal was therefore concentrated on these degradative routes rather than distributed broadly across the secretory arm.

A protein that several diseases engage can matter to its process in two ways, by shortening the routes through it and by holding it together at all, and these analyses separate the two. Single-node and sequential in silico node removal reduced path efficiency in the reference network. Single-node removal never changed the largest-component fraction, since no candidate is an articulation point, so the efficiency results are not corroborated by a separate connectivity outcome. Efficiency here is a property of paths in a graph, and the analysis does not measure substrate flux, processing rates or neuronal function. Here, a candidate shared network vulnerability is a protein among the largest structural contributions to cross-disease overlap whose in silico removal also produced a reduction in module path efficiency beyond the rewired-network null. Of the five largest contributors, PRKN, UBQLN2 and HSPA8 satisfy the efficiency criterion under the primary BH correction and represent the strongest candidate shared network vulnerabilities in the proteostasis network for cross-disease functional testing. None of the three satisfies both criteria under the conservative BY correction, so the pairing is a BH-level result and the three are put forward as candidates for experimental testing rather than as established vulnerabilities.

A small number of proteins recur across disease zones and account for a disproportionate share of the cross-disease structure, and the comparative map ranks them by that contribution in a way single-disease analyses cannot. PRKN, VCP, UBQLN2, ATXN3L and HSPA8 produced the largest structural contributions, although the neuronal functions and disease relevance of ATXN3L remain less established than those of the others. Their known biology is consistent with the structural roles observed here. In mitochondrial quality control, PRKN acts with PINK1 in a shared mitophagy pathway,^23–24,38^ and PINK1 itself recurred in seven of twelve primary disease zones and in eleven of the sixteen seed-protein neighborhoods, where its recurrence exceeded the matched null; its own structural contribution was not significant (*I*_*v*_ = 0.126; BH *q* = 1.00). In ubiquitin-dependent substrate handling, VCP performs substrate segregation and autophagosome maturation and is mutated in familial ALS/FTD,^25,39^ while UBQLN2 delivers ubiquitinated substrates to the proteasome and is mutated in X-linked ALS/FTD.^26^ CHMP2B was also a significant structural contributor. Its established role as an ESCRT-III component and its mutation in familial frontotemporal dementia^40^ connect the shared trafficking signal to endolysosomal membrane remodeling. PARK7 recurred in seven zones and contributed significantly to cross-disease overlap. In chaperone- and autophagy-mediated clearance, HSP70 chaperones triage misfolded substrates and suppress their toxicity in vivo.^27^ The constitutive chaperone HSPA8 delivers substrates such as alpha-synuclein to the lysosome through chaperone-mediated autophagy.^41^ The autophagy receptor SQSTM1 links ubiquitinated aggregates to autophagic clearance.^42^

Individually, each of these is a plausibility check. Together they describe the pattern a component-level architecture predicts, because convergence carried by the identity of individual components, rather than by membership of a process, requires single components to act on disease proteins beyond the disorder they are genetically linked to. Observations of that shape are already reported, one component at a time. The chaperone and chaperone-mediated autophagy route described above handles alpha-synuclein, and SQSTM1 also protects against huntingtin-induced cell death.^42^ Ataxin-2, the SCA2 protein, modifies TDP-43 proteinopathy in genetic and therapeutic experiments.^43–44^ VCP and UBQLN2 carry independent mutations across the ALS and frontotemporal dementia spectrum while performing general ubiquitin-dependent substrate handling.^25–26,39^ None of these observations was made to test a cross-disease architecture, and their agreement with it is independent of the present analysis.

Carrying the overlap between diseases and carrying the connections within a process are different jobs, and the proteins that do them are overlapping but not identical. Among the five largest structural contributors, PRKN, HSPA8 and UBQLN2 are supported on both counts. PARK7 and AKT1 are significant on both as well, with structural contributions well below those three (0.124 and 0.051, against 0.239 to 0.458), and are not carried forward as primary candidates. VCP contributes significantly to overlap, although its in silico removal did not produce a significant efficiency change in the ubiquitin-processing module. SQSTM1 shows the reverse pattern, reducing autophagy-module efficiency and being called differentially abundant in Alzheimer’s disease and FTLD-TDP without belonging to the nine significant overlap contributors, and PINK1 and HSP90AA1 are among the proteins that support efficiency alone. Removal of ATXN3L or ACTB produced small significant changes opposite to a loss of efficiency, which cautions against reading contribution to overlap as evidence that a protein holds a network together and is a reminder that removing a node also changes the graph over which efficiency is measured. Experimental evaluation should prioritize proteins whose structural contributions remain directionally consistent as evidence is restricted, and PRKN, VCP and HSPA8 retain positive effects across all four restricted regimes (Figure S3G).

Any analysis built on an association network faces a choice between breadth, which admits associations supported mainly by literature co-occurrence, and stringency, which discards real interactions that have not been assayed directly. Neither extreme is defensible alone, and the five evidence regimes lie between them, with the primary claims required to hold across them rather than at a single threshold. Individual pairwise relationships carry less weight than the global architecture, because global similarity exceeded its null under every evidence regime whereas the number of supported pairs fell as the confidence threshold rose. The supported sets were nested rather than reshuffled, and the relationships persisting under restricted evidence therefore form an ordered subset rather than a different answer. Alzheimer’s disease with Parkinson’s disease and Huntington’s disease with Parkinson’s disease are the cross-class members of that subset, each supported under three of the four restricted regimes. Together, the Alzheimer’s disease-Parkinson’s disease and Huntington’s disease-Parkinson’s disease relationships form the most evidence-robust cross-class component of the pairwise analysis. The ALS/FTD-SCA2 relationship is supported under two restricted regimes and is discussed here for its genetic and experimental context rather than for its evidence robustness, since intermediate-length ATXN2 expansions increase ALS risk and modify TDP-43 toxicity, and lowering ATXN2 reduces TDP-43 pathology and extends survival in a TDP-43 mouse model.^43–44^ The Parkinson’s disease and prion disease relationship concerns where the seed-protein neighborhoods of these diseases fall on a shared network. It is a statement about shared clearance architecture, distinct from the prion-like propagation of the disease-associated proteins themselves^34^ and not evidence of cross-seeding between alpha-synuclein and the prion protein. Where the diseases differ, they differ in emphasis rather than in inventory.

Divergence is carried by how they weight shared processes and not by robust sets of disease-unique proteins, and only SCA3 showed a primary-regime excess of single-zone proteins once zone size, proteostasis composition and degree were controlled. That excess is directionally consistent with the deubiquitinase activity of ATXN3^37^ although it is threshold-dependent, does not survive conservative correction, and remains a hypothesis rather than a disease-defining feature.

The PanNDA analysis showed that network position and molecular alteration provide different forms of evidence. Proteins in the Alzheimer’s disease zone underwent greater remodeling in the insoluble brain proteome, providing disease-matched support for an association between the network-defined APP-MAPT neighborhood and molecular change in affected human tissue. The whole-proteome association was directionally consistent but did not survive removal of the alpha-tubulin family from the zone, whereas the insoluble-fraction association survived both that removal and the removal of the three most influential zone proteins. The same models showed no zone association in schizophrenia or bipolar-disorder brain measured on the same platform, so zone membership does not simply mark proteins that shift in any diseased brain, although no insoluble psychiatric proteome exists to test that layer. Across disorders, alteration magnitude was not detectably associated with within-module influence or disease-zone recurrence. Structurally prioritized proteins may therefore complement candidates selected through differential abundance. Several qualifications bound this reading. The corresponding zone estimates in Parkinson’s disease and FTLD-TDP did not pass correction, which does not demonstrate a difference between diseases, and the influence estimate changed direction after conditioning on degree. Whether highly connected, functionally constrained proteins are held under tighter regulation and buffered against perturbation^45,46^ could not be resolved here, because the influence-specific association was not separable from connectivity, and the absence of a consistent differential-abundance call among central proteins is not evidence of buffering. The same reasoning cautions against reading co-expression hub status as molecular vulnerability,^47^ since co-expression hubs are defined from the disease measurements themselves whereas the structural influence used here is defined on a disease-independent interactome. Disease-zone membership was also not enriched among top-decile influence proteins. One reading of the dissociation is worth stating as a hypothesis rather than as a result. Network position is a property of the constitutive interactome and describes which components a disease protein engages, whereas differential abundance in end-stage post-mortem tissue describes how the cells that survived responded. Under that reading the two are not competing estimates of one quantity, and a dissociation between them is compatible with the architecture rather than evidence against it. The reading is testable, because position should track engagement, measured as interaction and as substrate handling, rather than abundance, and tissue sampled before end-stage neuronal loss would separate the two.

Three classes of experiment follow. Cross-disease perturbation experiments could determine whether controlled modulation of the candidate shared network vulnerabilities PRKN, UBQLN2 and HSPA8, and of VCP, which contributes as strongly to overlap but without a detectable efficiency effect, alters the handling of disease-associated proteins from more than one disease. Process-aligned experiments could evaluate autophagy or the ubiquitin-proteasome and protein-repair process in disease models that preferentially engage the corresponding process. Disease-preferential experiments could test the exploratory ATXN3-associated ubiquitin-processing neighborhood. The interpretation makes a falsifiable prediction, because proteins acting as shared network vulnerabilities should alter the handling of substrates from more than one disease when perturbed, and a consistent failure to observe this would argue against it. The analysis identifies network-structural sensitivities without establishing the direction or magnitude of any intervention, and whether modulating these proteins improves proteostasis function and neuronal resilience remains an unanswered experimental question.

## Limitations of the study

This study describes association structure and network-structural sensitivity; it does not establish biological intervention or its direction. STRING combined scores quantify confidence in functional or physical association, not pathway activity, effect size, expression, molecular flux, activation, inhibition or causal influence; module representation does not measure autophagic flux or proteasome activity; and the protein, module and topology perturbations are structural counterfactuals and graph perturbations, not cellular knockouts, treatments or rescues. Disease zones are built from the interaction neighborhoods of the seed proteins and inherit the coverage and study bias of the underlying association network, which is the reason individual pairwise relationships attenuate at stricter evidence thresholds and the reason the durable claims are based on the regime-robust global architecture. The NONTEXT regime requires each retained association to carry a non-text-mining evidence channel but does not recompute the STRING combined score with that channel removed, which means it bounds rather than eliminates the contribution of literature-derived evidence. Of the 1,045 proteins the regime retains from the 1,173 primary seed-neighborhood proteins, 66 carry no text-mining evidence and 567 are supported by an association for which text-mining is the strongest single channel; 491 of the 1,173 are retained when the combined score is recomputed from the non-text-mining channels alone (Table S23). Zone sizes vary widely, and the smallest zone (DRPLA) is low-coverage and descriptive only. A genetic comparison based on ALS association would address only the ALS component of the combined ALS/FTD zone. The ATXN3-associated ubiquitin-processing neighborhood is threshold-dependent and exploratory. The twelve diseases and sixteen seed proteins analyzed here are a defined set of diseases, and single-zone status is a statement about this set, not a claim of biological disease-specificity.

The molecular arm of this study also has specific limits. PanNDA comprises six neurodegenerative disorders, of which three (Alzheimer’s disease, Parkinson’s disease and FTLD-TDP) align directly with disease zones defined here; Huntington’s disease, prion disease and the polyglutamine ataxias are not represented, which limits the architecture-to-molecular comparison to three of the twelve diseases and leaves the cross-class relationships as network predictions for the remainder. The proteomic measurements are observational and derive from end-stage post-mortem tissue. Because the published statistics are already adjusted for the proportions of the five major brain cell types, an apparent lack of differential abundance among central proteins is less likely to be attributable to a shift in cell-type composition, but it cannot be fully separated from selective loss of the most affected neurons within a cell type; the disease-zone association, while robust to permutation, is modest in size, and the influence-specific associations were not separable from connectivity. We therefore frame the absence of a robust association between network position and molecular state as an observation to be tested functionally rather than as evidence of a protective mechanism. The whole-proteome disease-zone association also depends on the alpha-tubulin paralogues the zone inherits from MAPT and does not survive their removal, whereas the insoluble-fraction association does (Table S17).

## Methods

### Key resources table

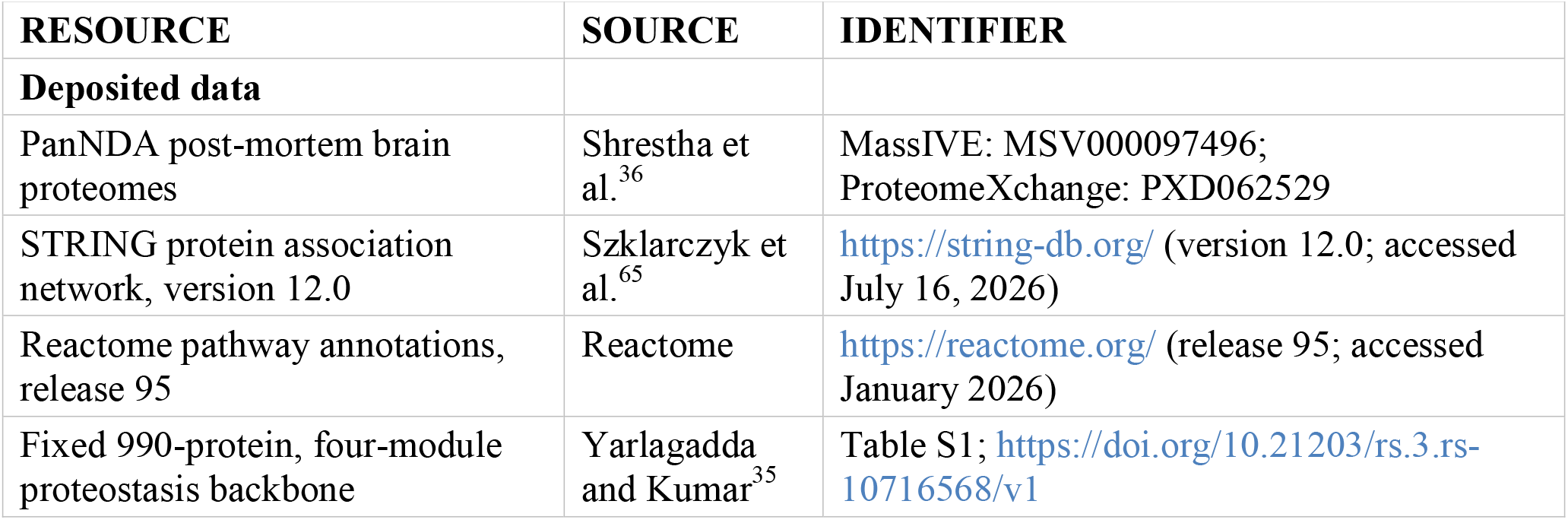
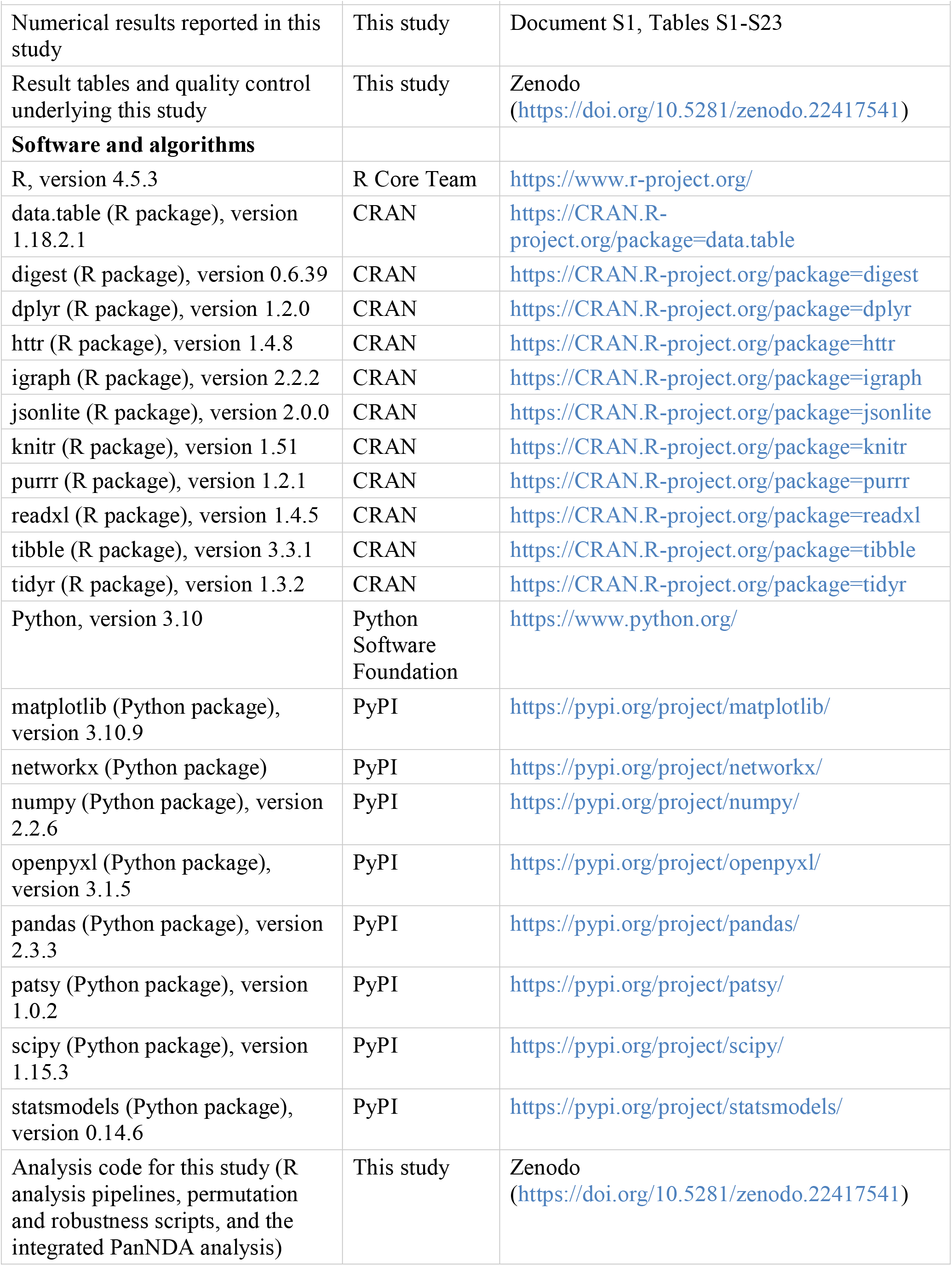

| RESOURCE | SOURCE | IDENTIFIER |
| --- | --- | --- |
| **Deposited data** |  |  |
| PanNDA post-mortem brain proteomes | Shrestha et al.<sup>36</sup> | MassIVE: MSV000097496; ProteomeXchange: PXD062529 |
| STRING protein association network, version 12.0 | Szklarczyk et al.<sup>65</sup> | https://string-db.org/ (version 12.0; accessed July 16, 2026) |
| Reactome pathway annotations, release 95 | Reactome | https://reactome.org/ (release 95; accessed January 2026) |
| Fixed 990-protein, four-module proteostasis backbone | Yarlagadda and Kumar<sup>35</sup> | Table S1; https://doi.org/10.21203/rs.3.rs-10716568/v1 |
| Numerical results reported in this study | This study | Document S1, Tables S1-S23 |
| Result tables and quality control underlying this study | This study | Zenodo<br>(https://doi.org/10.5281/zenodo.22417541) |
| **Software and algorithms** |  |  |
| R, version 4.5.3 | R Core Team | https://www.r-project.org/ |
| data.table (R package), version 1.18.2.1 | CRAN | https://CRAN.R-project.org/package=data.table |
| digest (R package), version 0.6.39 | CRAN | https://CRAN.R-project.org/package=digest |
| dplyr (R package), version 1.2.0 | CRAN | https://CRAN.R-project.org/package=dplyr |
| httr (R package), version 1.4.8 | CRAN | https://CRAN.R-project.org/package=httr |
| igraph (R package), version 2.2.2 | CRAN | https://CRAN.R-project.org/package=igraph |
| jsonlite (R package), version 2.0.0 | CRAN | https://CRAN.R-project.org/package=jsonlite |
| knitr (R package), version 1.51 | CRAN | https://CRAN.R-project.org/package=knitr |
| purrr (R package), version 1.2.1 | CRAN | https://CRAN.R-project.org/package=purrr |
| readxl (R package), version 1.4.5 | CRAN | https://CRAN.R-project.org/package=readxl |
| tibble (R package), version 3.3.1 | CRAN | https://CRAN.R-project.org/package=tibble |
| tidyr (R package), version 1.3.2 | CRAN | https://CRAN.R-project.org/package=tidyr |
| Python, version 3.10 | Python Software Foundation | https://www.python.org/ |
| matplotlib (Python package), version 3.10.9 | PyPI | https://pypi.org/project/matplotlib/ |
| networkx (Python package) | PyPI | https://pypi.org/project/networkx/ |
| numpy (Python package), version 2.2.6 | PyPI | https://pypi.org/project/numpy/ |
| openpyxl (Python package), version 3.1.5 | PyPI | https://pypi.org/project/openpyxl/ |
| pandas (Python package), version 2.3.3 | PyPI | https://pypi.org/project/pandas/ |
| patsy (Python package), version 1.0.2 | PyPI | https://pypi.org/project/patsy/ |
| scipy (Python package), version 1.15.3 | PyPI | https://pypi.org/project/scipy/ |
| statsmodels (Python package), version 0.14.6 | PyPI | https://pypi.org/project/statsmodels/ |
| Analysis code for this study (R analysis pipelines, permutation and robustness scripts, and the integrated PanNDA analysis) | This study | Zenodo<br>(https://doi.org/10.5281/zenodo.22417541) |

### Resource availability

#### Lead contact

Further information and requests for resources should be directed to the lead contact, Vijay Kumar.

#### Materials availability

This study did not generate new unique reagents.

#### Data and code availability

All reported numerical results are provided in the manuscript and Supplemental Tables S1-S23. All custom analysis code used in this study, including the R Markdown analysis pipelines, the permutation and robustness testing scripts, the Python architecture-state analysis, and a record of the computational environment (R, Python and package versions), is archived and publicly available on Zenodo (https://doi.org/10.5281/zenodo.22417541). The same archive carries the input data for the network analyses, namely the STRING v12.0 seed-module snapshots, the fixed APP/MAPT reference files, and the Reactome-derived module assignment, together with this study’s derived result tables; STRING data are redistributed under the Creative Commons Attribution 4.0 International license that resource applies to its data. Every analysis other than the PanNDA arm can therefore be rerun from the archive alone. The PanNDA per-protein statistics are the original study’s supplementary material and are not redistributed here; the archive documents how to obtain the source workbooks. Any additional information required to reproduce the results is available from the lead contact. The proteomic data analyzed here were generated by the original PanNDA study^36^ and are available via MassIVE (MSV000097496) and ProteomeXchange (PXD062529); processed per-protein statistics are in the supplemental tables of that publication.

### Method details

#### Disease-naive proteostasis framework and eligible universe

A fixed, disease-naive proteostasis backbone of 990 proteins was used, constructed as described previously.^35^ In brief, proteins representing the principal systems of protein quality control were curated from Reactome pathway annotations (release 95, accessed January 2026) and partitioned, on the basis of their principal annotated function, into four mutually exclusive modules (M1 vesicular trafficking; M2 autophagy; M3 ubiquitin-proteasome system and protein repair; M4 chaperones and folding); module boundaries were checked for functional coherence against Gene Ontology, KEGG and Reactome gene-set enrichment. The curated set was mapped to STRING v12.0, and only associations among the curated proteins were retained, with no external interactors added, at a combined score of at least 0.4. Reactome annotated 996 proteins across the four modules, of which 75 were annotated in more than one module and were assigned a single primary module by the function-based rule above. Six proteins were then removed, three excluded before the STRING mapping and three retaining no within-module association at the primary threshold, giving the fixed 990-protein backbone (M1 vesicular trafficking, 478; M2 autophagy, 102; M3 ubiquitin-proteasome system and protein repair, 331; M4 chaperones and folding, 79); the per-protein record is deposited as 00_node_attrition_table.csv. The framework was assembled before any disease data were introduced. Sixteen disease-associated seed proteins were mapped to twelve diseases (Alzheimer’s disease: APP, MAPT; ALS/FTD: TARDBP, FUS, SOD1, C9orf72; Parkinson’s disease: SNCA; prion disease: PRNP; Huntington’s disease: HTT; and the polyglutamine disorders SCA1 [ATXN1], SCA2 [ATXN2], SCA3 [ATXN3], SCA7 [ATXN7], DRPLA [ATN1], SBMA [AR] and SCA17 [TBP])^2–8,48–56^. Four disease-associated seed proteins (APP, AR, ATXN3, ATXN7) are backbone residents and were removed from every eligibility-restricted analysis, leaving 986 eligible proteins.

#### Selection of diseases and their disease-associated seed proteins

The disease panel and its disease-associated seed proteins were fixed in the locked prespecification before any comparative testing. The panel comprises twelve diseases represented by sixteen disease-associated seed proteins. Each disease-protein relationship and its class assignment were taken from the membership table (04_disease_zone_membership.csv) and are keyed to the identifiers in that table. Because the backbone is outcome-independent and was assembled before disease data were introduced, every disease-associated seed protein is mapped onto the same reference network by the identical seed-neighborhood rule, and the two diseases with more than one disease-associated seed protein (Alzheimer’s disease and ALS/FTD) are handled by the prespecified within-disease intersection. Single-zone status is therefore a property of this fixed, finite disease panel and does not establish biological disease specificity or generalizability beyond the diseases examined. The panel was assembled by literature-informed curation rather than by a systematic database screen. A disease was included when it was an established neurodegenerative proteinopathy represented by one or more proteins with a well-established defining pathological association or encoded by a causal disease gene, and could be mapped onto the fixed proteostasis backbone through STRING v12.0. Alzheimer’s disease and ALS/FTD were represented by more than one protein because more than one disease-associated seed protein is well established for each. The eight polyglutamine disorders were included as a genetically coherent repeat-expansion family that nonetheless acts through distinct proteins, supporting both within-family and cross-family comparison. Conditions lacking a prespecified protein with a well-established defining pathological association or causal genetic link, or for which the selected protein could not be mapped consistently onto the backbone, were not included, which ensured that every disease-to-protein assignment was prespecified, literature-supported and outcome-independent.

#### Disease-zone construction and evidence regimes

For disease-associated seed protein g, module m and regime r, the seed neighborhood is 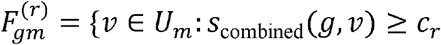 and *Q*_*r*_ (*g, v*)}, where *U*_*m*_ is the module protein set, *s*_combined_ is the STRING combined score and *Q*_*r*_ is the regime evidence predicate. Here *G*_*d*_ is the set of disease-associated seed proteins of disease *d*. The disease zone is the union across modules of the proteins shared by every disease-associated seed protein within that module, 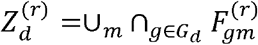. Disease-associated seed proteins resident in the backbone are then removed to give the eligible zone used for all primary analyses, 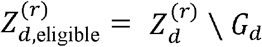. Five regimes were evaluated: C400 (combined score ≥ 0.4, primary), C700 (≥ 0.7), C900 (≥ 0.9), a regime requiring non-text-mining support (NONTEXT; ≥ 0.4 with at least one non-text-mining evidence channel), and an experimental- or database-supported regime (EXPDB; ≥ 0.4 with experimental or database evidence). A protein-protein association is retained under a regime when any raw evidence row for that pair satisfies the complete regime predicate.

#### Similarity and global statistic

Pairwise similarity is the Jaccard index *J*(*A, B*) = |*A* ∩ *B*|/|*A* ∪ *B*|. The global statistic is *T*_global_ = ∑_1≤ *i*<*j*≤12_ *J* (*Z*_*i*_, *Z*_*j*_), the summed similarity over all 66 disease pairs.

#### Polyglutamine-family robustness

Because the panel contains eight polyglutamine disorders (Huntington’s disease and the seven repeat-expansion ataxias and atrophies; 28 of the 66 disease pairs), we assessed whether this global overlap depended on this family under the same C400 joint null (100,000 permutations; module-by-within-module-degree-bin stratification, overlap allowed across diseases). We computed a cross-class statistic restricted to the 38 disease pairs that are not internal to the polyglutamine family; the decomposition of the global statistic into its cross-class and within-polyglutamine contributions; leave-one-disease-out statistics, recomputing the summed pairwise Jaccard over the remaining eleven diseases; and family-balanced five-disease panels comprising the four non-polyglutamine diseases and one polyglutamine representative, evaluated for each of the eight representatives. Each statistic used add-one upper-tail permutation p-values under the same null (Table S5).

#### Within-disease overlap

For the two diseases with more than one disease-associated seed protein, the number of proteostasis proteins shared across their seed-protein neighborhoods was compared with a composition-matched null (10,000 permutations); a cross-protein recurrence analysis additionally counted, for each protein, the number of the sixteen seed-protein neighborhoods containing it, evaluated against the same null.

#### Module-dependence statistic

With *n*_*db*_ the protein count of the zone in global degree bin b, *Nb* the eligible-backbone count in bin b and *N*_*bm*_ the eligible module-m count in bin b, the expected module count is *E*_*dm*_ = ∑_*b*_ *n*_*db*_ (*N*_*bm*_/*N*_*b*_). The per-module component is *C*_*dm*_ = (*O*_*dm*_ − *E*_*dm*_)^2^/ *E*_*dm*_, the omnibus statistic is *T*_*d*_ = ∑_*m*_ *C*_*dm*_, and the direction of a combination is sign(*O*_*dm*_ − *E*_*dm*_). Global degree bins were defined once from the fixed degree table by ascending degree then alphabetical symbol into ten equal bins; a combination with *E*_*dm*_ = 0 is reported as not testable.

#### Structural perturbations

Protein ablation removes a protein from every zone: 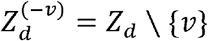; 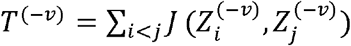; and the structural contribution of the protein is *I*_*v*_= *T* − *T*^(−*v*)^, computed for all 986 eligible proteins by a closed-form algebraic identity verified against brute-force deletion, during the same C400 permutations used for the global analysis. Module ablation removes the proteins of a module: 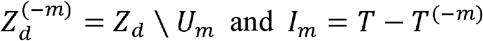. Topology knockout removes a single node from the simple undirected fixed-confidence graph of a module; efficiency is 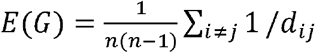, with unreachable pairs contributing zero; the largest-component fraction is *L*(*G*) = | *C*_max_(*G*)|/| *V*(*G*)|; and single-node effects are Δ*Ev*= *E* (*G*) − *E* (*G* −*v*) and Δ*L*_*v*_= *L*(*G*) − (*G* −*v*). Candidate protein sets comprised proteins present in at least six disease zones and the single-zone proteins of the disease with the strongest primary single-zone signal. Robustness curves removed nodes sequentially at fractions {0,0.01,0.025,0.05,0.10,0.15,0.20} ranked by recurrence, by within-module influence, and by degree (descriptive), and compared the trapezoidal area under *E* (*G*_*k*_)/ *E* (*G*_0_) and *L*(*G*_*k*_)/*L* (*G*_0_) with a matched-random null for the two inferential rankings.

#### Integrated Value of Influence

The Integrated Value of Influence (IVI), called within-module influence in the text, was computed within each module as IVI = Hubness × Spreading, with Hubness = *DC*_*sc*_ + *LH*_*sc*_ and Spreading = (*NC*_*sc* +_ *CR*_*sc*_)(*BC*_*sc*_ + *CI*_*sc*_), from minmax-scaled degree, local h-index, neighborhood connectivity, ClusterRank, weighted betweenness and collective influence, and interpreted within modules and not across them.

#### Topological clustering of disease zones

For each disease and module, the topological autocorrelation of binary zone membership over the simple undirected interaction graph of the module was quantified by the Moran statistic, 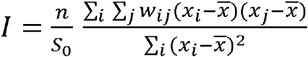, where w_*ij*_ is the row-standardized adjacency weight of the module graph, *x*_*i*_ is zone membership, 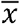 the mean membership and *S*_0_ = ∑_*i*,*j*_ *w*_*ij*_. Significance was assessed against 5,000 within-module membership-preserving permutations, with BH correction across the 36 testable disease-module combinations.

#### Null models and inference

The global and pairwise analyses and the single-zone analysis used a joint null that randomizes each zone within its module-by-degree-bin counts, with overlap allowed across diseases. Module dependence used a global-degree-bin null that does not preserve module membership. A neighborhood-reconstructed sensitivity randomized seed neighborhoods within their module-by-degree-bin counts and re-intersected them within multi-protein diseases. Topology knockouts used degree-sequence-preserving (Maslov-Sneppen) edge-switching rewirings of each module graph. Robustness curves used matched-random removal sequences drawn from the same module and within-module-degree bin as the corresponding targeted protein. Empirical p-values were add-one permutation values *P* = (*K* + 1)/(*B* + 1), reported with exact Clopper-Pearson intervals alongside K and B; *B* = 100,000 for the evidence-regime analyses. The similarity, composition (upper-tail on the non-negative module-combination statistic, with the direction of each taken from the signed observed-minus-expected deviation, such that the squared statistic captures departures in either biological direction), recurrence, and protein- and module-ablation contribution tests used upper-tail p-values; the single-node topology-deletion tests (network efficiency and largest-component change) used two-sided p-values, because network-structural ablation of a node may either increase or decrease these quantities; the robustness area-under-curve tests used lower-tail p-values, because a smaller area indicates faster network degradation; and the topological-clustering analysis used a two-sided permutation p-value, *p*_two-sided_ = min [1, 2min(*p*_greater_, *p*_less_)]. BH was the primary correction and BY a conservative sensitivity, each computed within fixed, non-pooled families. Any result whose BH q fell in the interval 0.04,0.06] was re-run as a full independent permutation sequence under a second seed; the primary single-zone result and the neighborhood-reconstructed sensitivity were always independently replicated.

#### Method references

Add-one permutation p-values and their bounds;^57–58^ multiple-testing correction;^59–60^ degree-preserving rewiring;^61^ the Integrated Value of Influence;^62^ spatial autocorrelation;^63^ exact binomial intervals;^64^ and the STRING association network and its evidence channels.^65^ The constructions of *T*_global_, *E*_*dm*_, *C*_*dm*_ and *T*_*d*_ are specific to this study and are evaluated through the permutation nulls above.

#### Integrated analysis of PanNDA brain proteomes

To relate network position to molecular state, we analyzed the Pan-Neurodegeneration Atlas (PanNDA)^36^, post-mortem brain proteomes of six neurodegenerative disorders (mass spectrometry data MassIVE MSV000097496 / ProteomeXchange PXD062529; processed per-protein statistics from Supplemental Tables S2, S3 and S5 of that study, and pathology-correlation lists from that study’s Supplemental Table S6). In the original study, protein abundances were corrected for age, sex, post-mortem interval and batch by linear regression, and for the proportions of five major brain cell types by constrained regression, before differential testing by moderated *t*-test; the per-protein statistics analyzed here are therefore already adjusted for those factors. PanNDA diseases were aligned to the present disease zones as Alzheimer’s disease, Parkinson’s disease and FTLD-TDP (aligned to ALS/FTD). From each disease statistical-analysis sheet we retained, per entity, the reported P value, false-discovery rate (FDR) and z-normalized log_2_ fold change, and reconstructed a uniform whole-proteome differentially-abundant-protein (DAP) call as FDR < 0.05 and |log_2_FC-z| > 2.5; under this harmonized rule the counts matched the published values for Alzheimer’s disease, Parkinson’s disease, Lewy body dementia (LBD) and FTLD-TDP, whereas progressive supranuclear palsy (PSP) gave 0 and vascular dementia (VAD) 9, against 248 and 296 under the disease-specific criterion the original study applied to those two cohorts (*P* < 0.01 with the same |log_2_FC-z| > 2.5 threshold); no PSP protein reached FDR < 0.05 in that cohort. Both diseases were retained in the harmonized six-disease binary and recurrence analyses under the uniform FDR-based criterion; data sources, provenance and differential-abundance extraction quality control are reported in Tables S19 and S20. Entities were mapped to the 986 analysis-eligible backbone proteins by UniProt accession first and gene symbol second; magnitude of alteration was the rank-inverse-normal transform of |log_2_FC-z| computed across the full measured proteome of each disease. Disease-zone engagement was tested per aligned disease, in both the whole-proteome and insoluble layers, by regressing magnitude on zone membership with module and within-module degree as covariates (heteroscedasticity-consistent HC3-robust linear models on continuous alteration magnitude) with a stratified label-permutation P value for the zone term.

Architecture-state coupling related alteration magnitude to within-module influence percentile and to disease-zone count in protein-by-disease models with module and disease fixed effects and protein-cluster-robust inference. Because within-module influence and within-module degree were strongly collinear (Spearman *ρ* = 0.83), the primary estimand was the total, degree-unconditional association, tested by a cluster-robust model and a matched degree-unconditional permutation, with a degree-adjusted permutation retained as a sensitivity analysis. Conditioning on within-module degree reverses the sign of the influence coefficient to *β* = +0.20 (permutation *P* = 0.010); because the estimate and its sign depend on the specification, neither the total nor the degree-conditional estimate is interpreted directionally (Table S17). Site-level differential-abundance calls in the phosphosite and ubiquitin-site layers used the original study’s criterion for post-translational-modification (PTM) enriched data, *P* < 0.05 with |log_2_FC-z| > 2.5, rather than the uniform FDR rule, which applies to the whole-proteome layer. Phosphosite- and ubiquitin-site-layer analyses used site-level models with parent-protein clustering and a parent-protein-level permutation in which all sites of a protein are permuted together. Cross-disease molecular recurrence was compared with disease-zone count by Spearman correlation over proteins with complete (six non-missing) whole-proteome DAP calls, the same complete-coverage set used for the four-state quadrant; proteins with any missing call were excluded rather than treated as unaltered. As a post-hoc disease control, the disease-zone models were refitted on the PsychENCODE schizophrenia and bipolar-disorder whole proteomes (Supplemental Table S5 of that study, sheets ST5.21 and ST5.20), under the same covariates, permutation strata, seed and permutation count; those cohorts carry no insoluble layer. Pathology-correlation lists were mapped onto the architecture descriptively, without enrichment tests, because the published lists are filtered to significant correlates and do not define a complete tested background. Permutation tests used 100,000 iterations within analysis-specific, non-pooled BH families. The stratification was not identical across tests: the null, permutation strata, tail and correction family of each analysis are stated individually in Table S22, and strata containing fewer than ten proteins were pooled within module. The binary differentially-abundant-protein secondary outcomes of the architecture-state analysis (stratified label permutation) and the whole-versus-insoluble interaction (Freedman-Lane permutation) were permutation-tested. The disease-zone models were also refit with zone-resident proteins removed, one at a time and then in the set of three whose removal moved the estimate furthest toward zero, under the same design, permutation scheme, seed and permutation count. Architecture-state results are reported in Table S17, and exploratory contributor and pathology analyses in Table S18. The analysis code and executed statistical specifications are archived on Zenodo (see data and code availability).

### Quantification and statistical analysis

The network analyses were performed in R version 4.5.3; the integrated PanNDA architecture-state analyses were performed in Python (statsmodels, scipy; seed 20260901, 100,000 permutations). Empirical p-values were calculated as p=(K+1)/(B+1). Global and pairwise similarity, proteostasis composition, single-zone and ablation analyses used 100,000 permutations in each STRING evidence regime. Within-disease overlap and cross-seed recurrence used 10,000 permutations; topological-knockout and robustness analyses used 10,000 permutations per process-specific network; and Moran’s I used 5,000 permutations per disease-process combination. Tail directions for every test, and the reasons for them, are specified under Null models and inference above, together with BH as the primary correction and BY as a conservative sensitivity within fixed, non-pooled families. Formal support for a disease pair required a significant global test and pairwise BH (*q* < 0.05). Monte Carlo precision, independent-replication results and the complete statistical specification are reported in Tables S21 and S22.

## Acknowledgments

The authors thank the developers of R, igraph and the STRING database for maintaining the open-source resources used in this analysis. The disease-naive proteostasis backbone analyzed here was defined in the associated single-disease study. We thank the investigators of the Pan-Neurodegeneration Atlas (PanNDA) for making their proteomic data publicly available. V.K. sincerely acknowledges Sharda University Agra for providing infrastructural facilities. This study received no external funding.

## Supplemental information

Supplemental information includes Figures S1-S17 and Document S1. Tables S1-S23.

