## Supplementary Figures for "Genetically distinct neurodegenerative proteinopathies engage overlapping proteostasis components"

Figures S1 to S17. Supplemental tables S1 to S23 are provided separately as Document S1.

**Figure S1. The 990-protein, four-module proteostasis network, related to Figures 2 and 3.** The fixed proteostasis backbone (990 proteins mapped to STRING v12.0, combined score at least 0.4) shown as its four biological-process modules: vesicular trafficking (M1, 478 proteins), autophagy (M2, 102), ubiquitin-proteasome processing and protein repair (M3, 331) and chaperones and protein folding (M4, 79). Node size scales with within-module degree and edges are the within-module STRING associations. Proteins present in at least six of the twelve disease zones are ringed and labeled. See also Table S1.

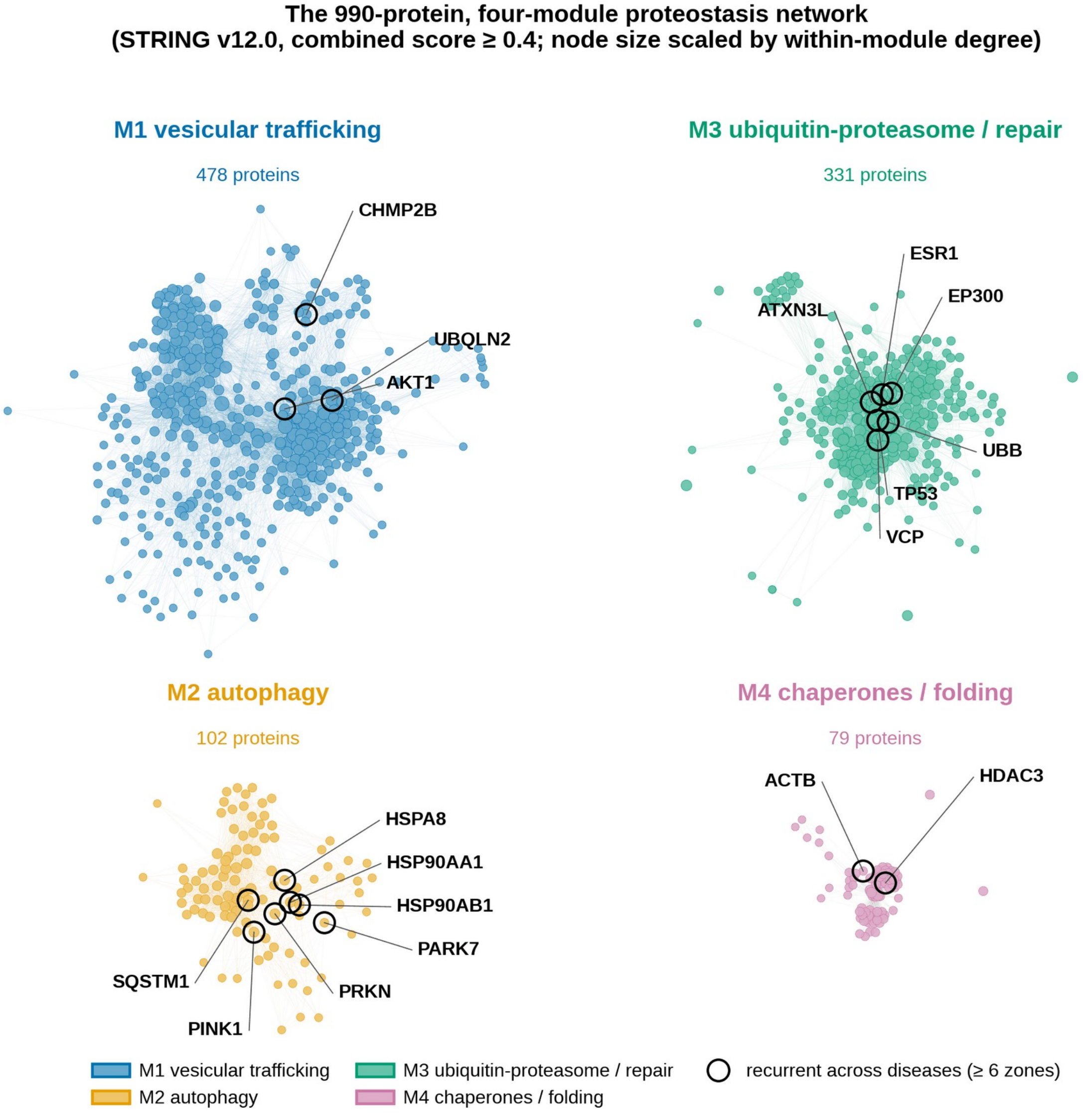

**Figure S2. Sensitivity, structural and disease-preferential analyses of the shared network, related to Figures 3-5.** (A) Global disease-zone similarity ($T_{\text{global}}$) against the null band for the primary analysis and three reconstructions (excluding DRPLA, retaining seeds, neighborhood-reconstructed). (B) Number of formally supported disease pairs under each. (C) Average-linkage similarity dendrogram on 1 - Jaccard. (D) Proteostasis-protein count of each disease zone in each process. (E) Disease-specific topological clustering (Moran I per disease and process; asterisks BH $q < 0.05$; gray, not testable). (F) Single-zone (disease-restricted) protein count per disease against the null band; only SCA3 exceeds its null after correction (*BH $q < 0.05$). See also Tables S4, S6, S8 and S16.

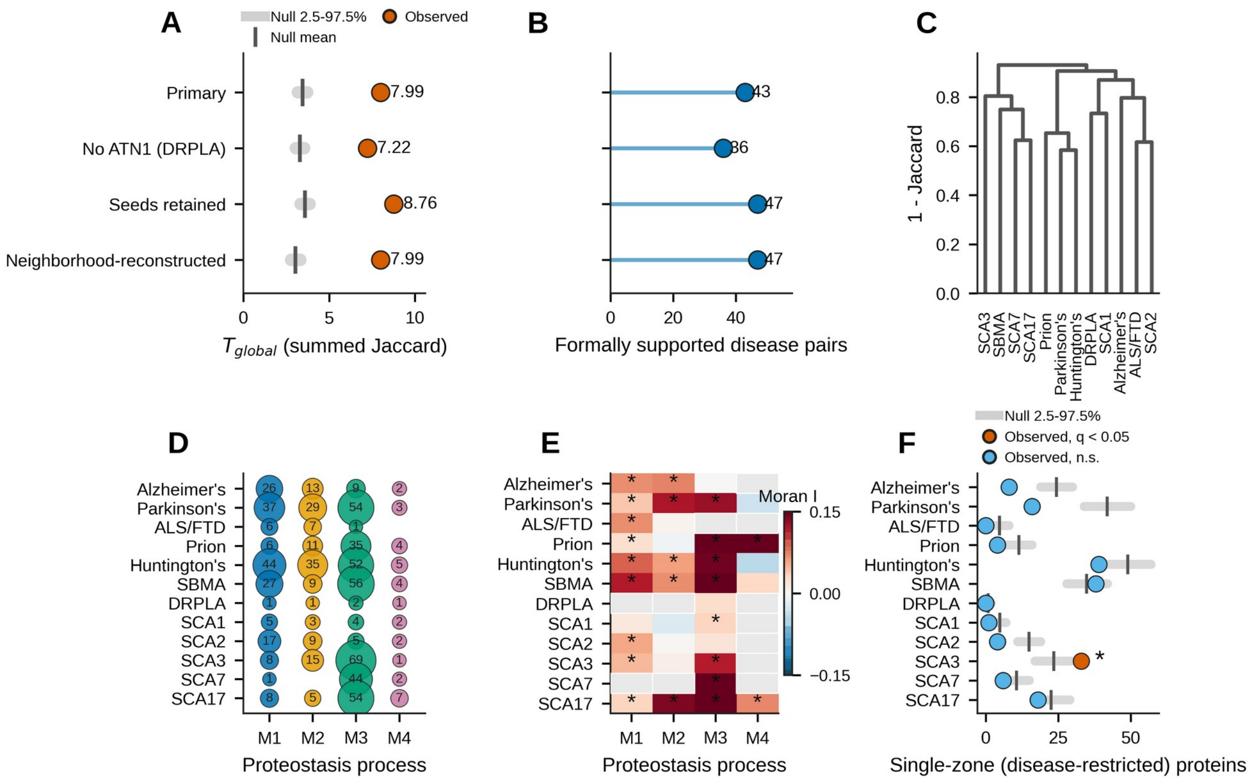


**Figure S3. Evidence-regime contraction, selected pairwise null comparisons, the exploratory ATXN3 neighborhood, and protein-ablation sensitivity, related to Figures 4, 6 and 7.** (A) Disease-zone size (mean and min-max range) contracts as STRING evidence stringency increases across the five regimes. (B) Observed versus null-mean Jaccard similarity for selected cross-class pairs. (C) Pairwise protein-ablation effect (change in Jaccard on removal) of leading proteins across selected disease pairs. (D) Exploratory ATXN3-associated ubiquitin-processing neighborhood in SCA3: observed single-zone count versus the matched-null mean across the five STRING evidence regimes; the excess is significant only at the C400 regime (asterisk, BH $q < 0.05$) and is reported as exploratory. (E) SCA3 single-zone significance under BH and the conservative BY correction across evidence regimes. (F) Change in network efficiency ($\Delta E_v$) on network-structural single-node ablation for the 33 ATXN3 single-zone candidates (vermilion, BH $q < 0.05$, 23 of 33; black-ringed, also BY $q < 0.05$, six; gray, not significant). (G) Direction of the twelve leading structural contributors (the nine with a significant structural contribution after correction, plus HSP90AA1, PINK1 and SQSTM1) across the four restricted evidence regimes (green, concordant with the primary regime). See also Tables S2, S4, S12, S13, S14 and S16.

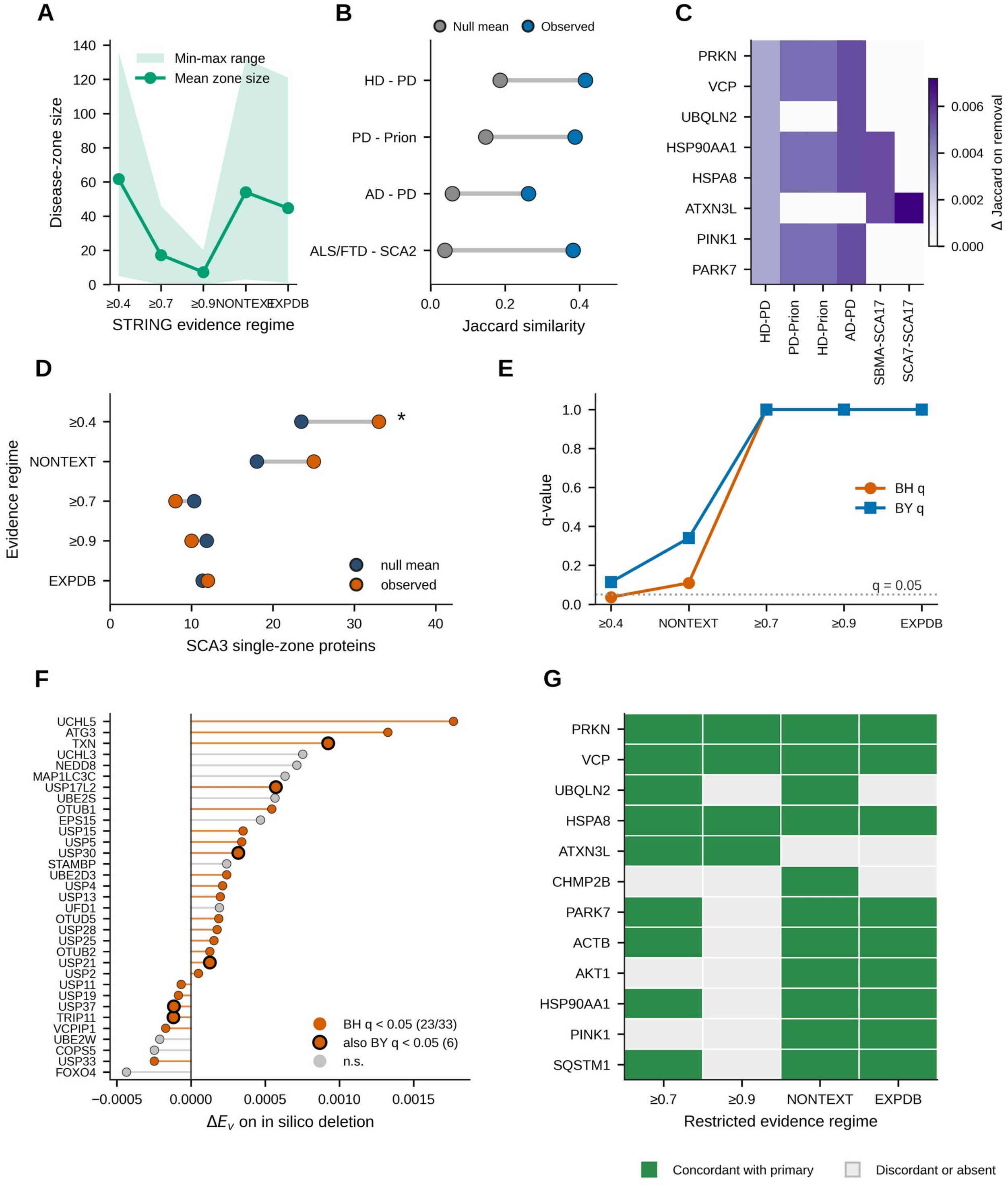

A
Min-max range
Mean zone size
Disease-zone size
STRING evidence regime
≥0.4
≥0.7
≥0.9
NONTEXT
EXPDB
B
Null mean
Observed
HD - PD
PD - Prion
AD - PD
ALS/FTD - SCA2
Jaccard similarity
C
PRKN
VCP
UBQLN2
HSP90AA1
HSPA8
ATXN3L
PINK1
PARK7
HD-PD
PD-Prion
HD-Prion
AD-PD
SBMA-SCA17
SCA7-SCA17
Δ Jaccard on removal
D
Evidence regime
null mean
observed
SCA3 single-zone proteins
E
q-value
BH q
BY q
q = 0.05
F
UCHL5
ATG3
TXN
UCHL3
NEDD8
MAP1LC3C
USP17L2
UBE2S
OTUB1
EPS15
USP15
USP5
USP30
STAMBP
UBE2D3
USP4
USP13
UFD1
OTUD5
USP28
USP25
OTUB2
USP21
USP2
USP11
USP19
USP37
TRIP11
VCPIP1
UBE2W
COPS5
USP33
FOXO4
BH q < 0.05 (23/33)
also BY q < 0.05 (6)
n.s.
ΔE_v on in silico deletion
G
PRKN
VCP
UBQLN2
HSPA8
ATXN3L
CHMP2B
PARK7
ACTB
AKT1
HSP90AA1
PINK1
SQSTM1
Restricted evidence regime
Concordant with primary
Discordant or absent

**Figure S4. Robustness of the shared architecture to STRING evidence regime (A, B) and to targeted node removal (C, D), related to Figures 4, 5 and 7.** (A) Pairwise disease-zone Jaccard similarity at each of the five evidence regimes. (B) Relative process composition per disease at each regime. (C) Efficiency ratio and (D) largest-component ratio against the fraction of nodes removed, for each process network under recurrence-, IVI- and degree-ranked removal. See also Tables S4, S7 and S15.

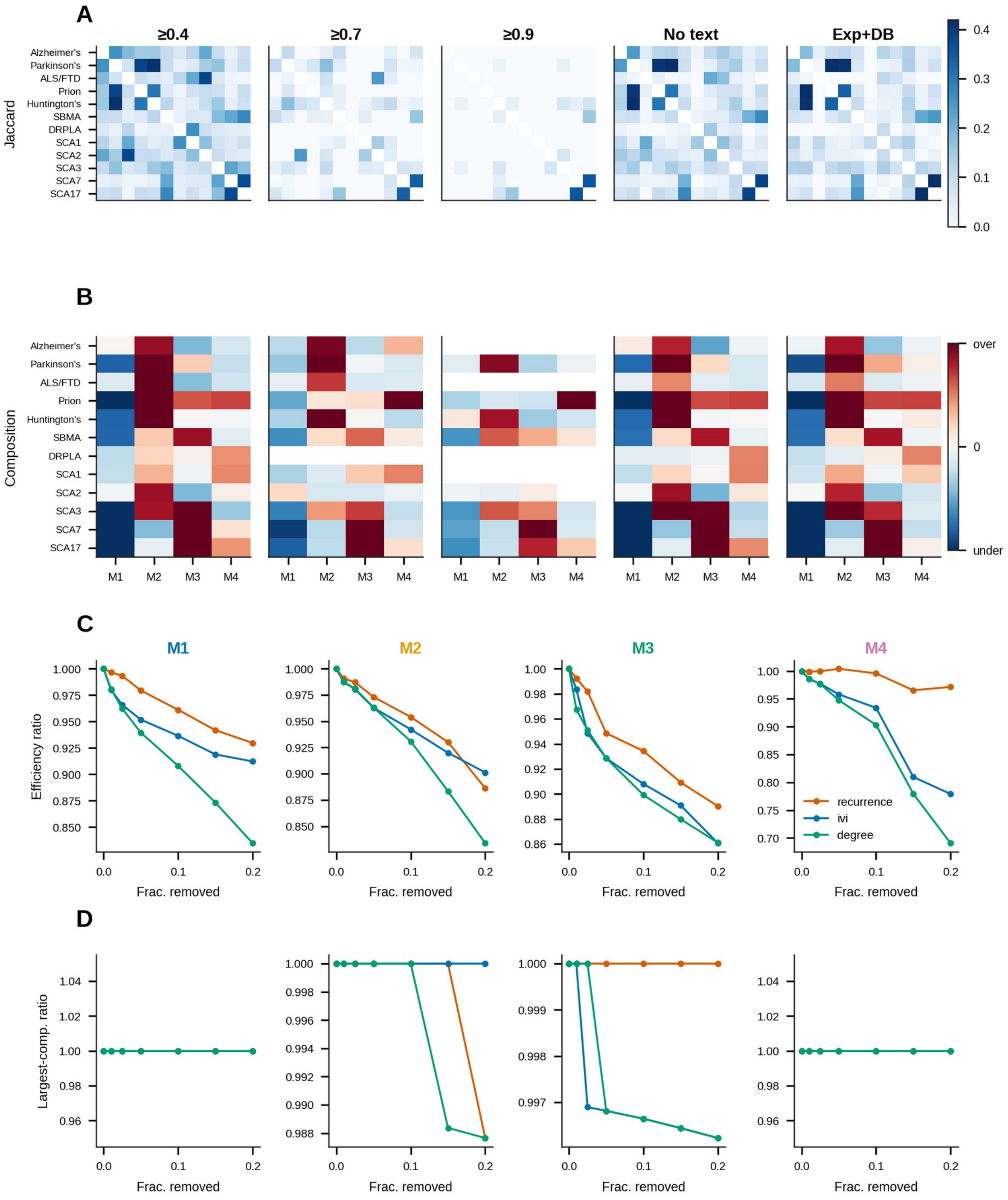

**Figure S5. Supplementary analyses for the integrated PanNDA arm, related to Figure 7.** (A) Whole versus insoluble signed alteration for Parkinson's disease backbone proteins; Parkinson's disease-zone proteins in orange, Pearson *R* shown. (B) The same comparison for FTLD-TDP. (C) Architecture-state estimates for within-module influence (IVI) and structural recurrence, comparing the primary total model, the degree-adjusted sensitivity model (which reverses sign and is unstable), and the binary differential-abundance model; permutation P shown. (D) Whole-minus-insoluble layer interaction for IVI and recurrence.

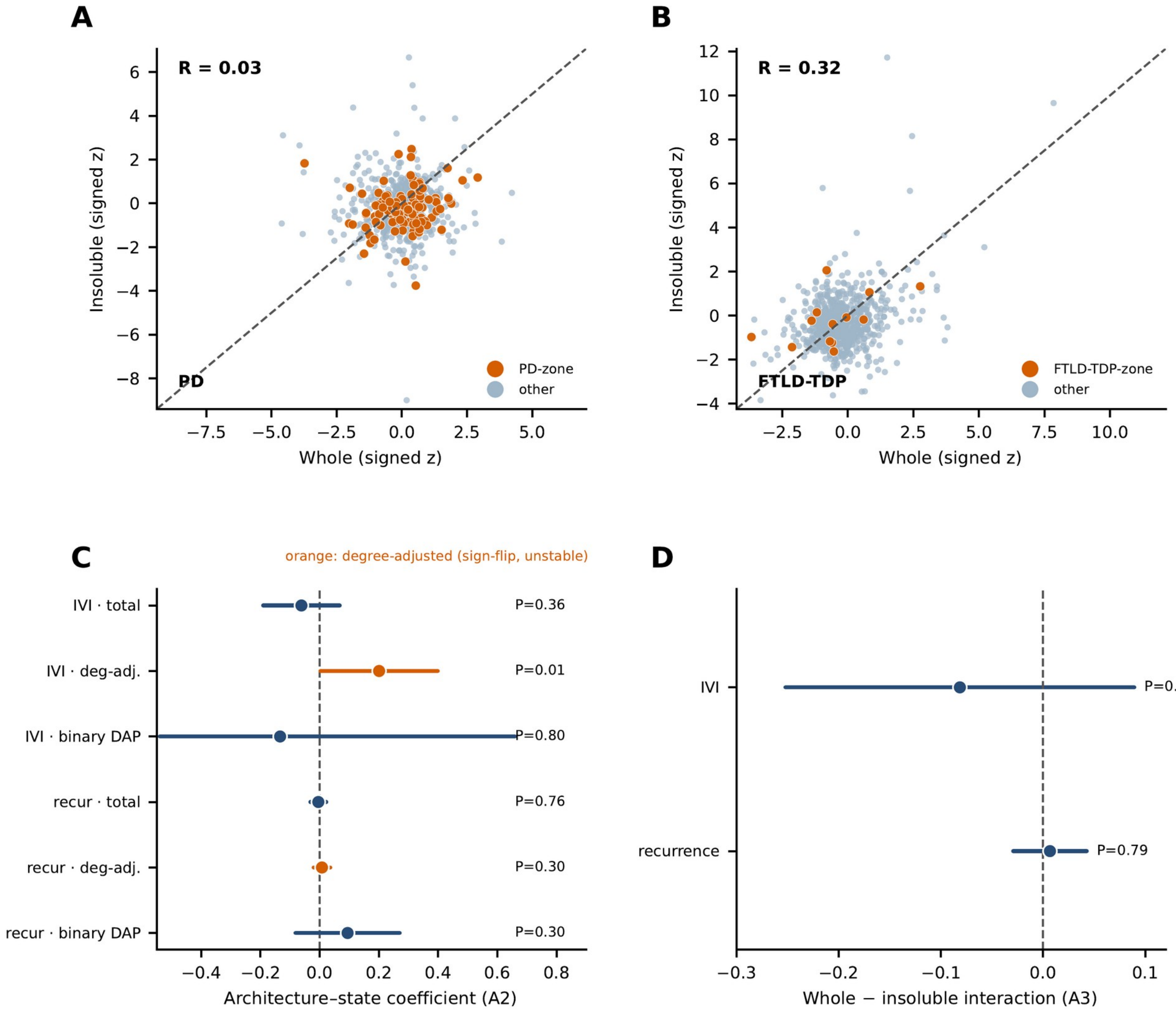

**Figure S6. Complete proteostasis neighborhood of Alzheimer's disease, related to Figures 2 and 3.** Every protein in the primary seed-removed Alzheimer's disease zone (C400; eligible universe; 50 proteins) is shown as a network: nodes are the zone proteins, grouped around the ring by biological process (vesicular trafficking M1, autophagy M2, ubiquitin-proteasome processing and protein repair M3, chaperones and protein folding M4) and colored accordingly, and chords denote within-zone STRING associations (combined score at least 0.4). Proteins present in at least six of the twelve disease zones are ringed and shown in bold; disease-restricted proteins (present in a single zone) are shown in gray. Disease-associated seed protein(s): APP, MAPT. See also Table S2.

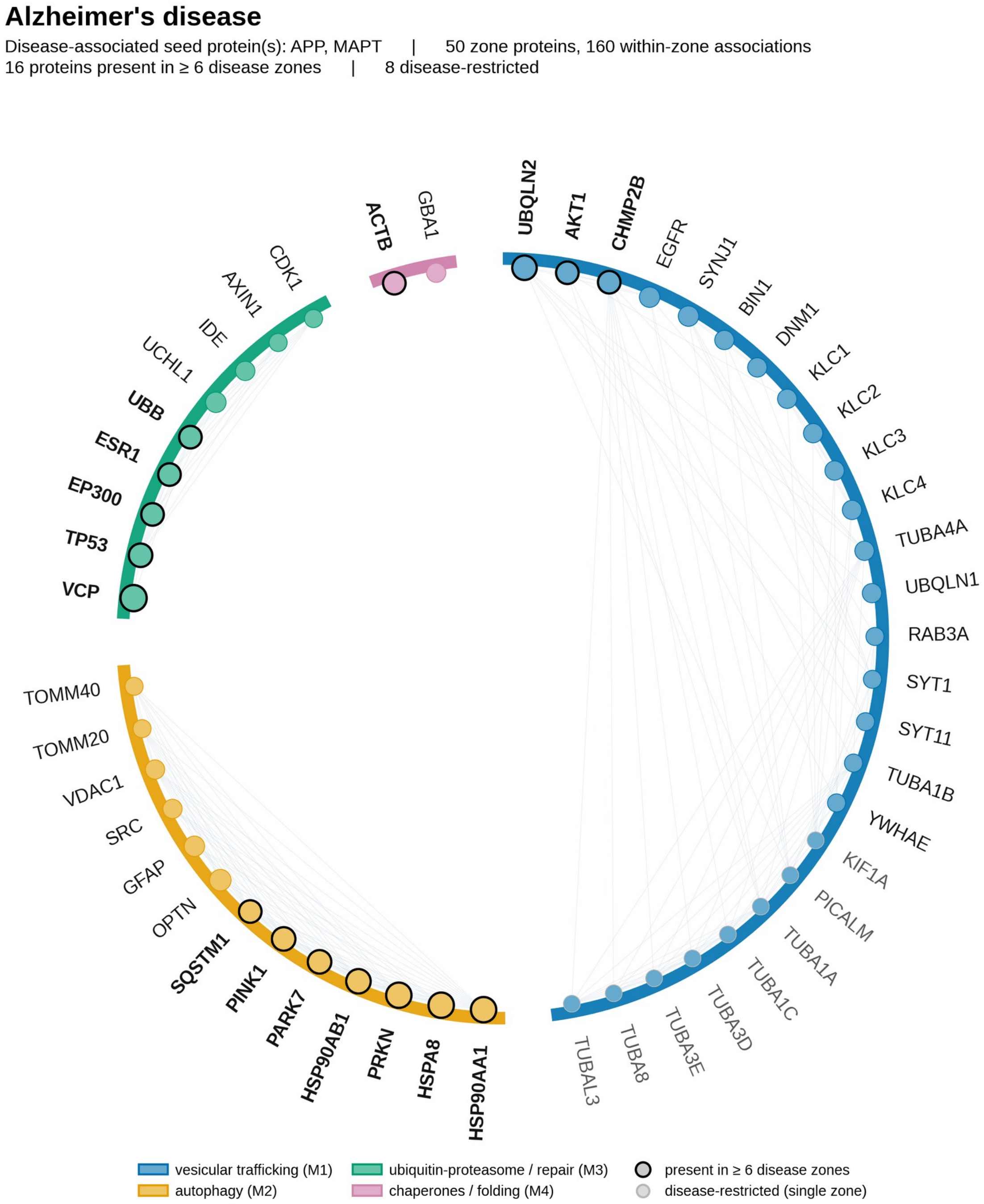

**Figure S7. Complete proteostasis neighborhood of ALS/FTD, related to Figures 2 and 3.** Every protein in the primary seed-removed ALS/FTD disease zone (C400; eligible universe; 14 proteins) is shown as a network: nodes are the zone proteins, grouped around the ring by biological process (vesicular trafficking M1, autophagy M2, ubiquitin-proteasome processing and protein repair M3, chaperones and protein folding M4) and colored accordingly, and chords denote within-zone STRING associations (combined score at least 0.4). Proteins present in at least six of the twelve disease zones are ringed and shown in bold; disease-restricted proteins (present in a single zone) are shown in gray. Disease-associated seed protein(s): TARDBP, FUS, SOD1, C9orf72. See also Table S2.

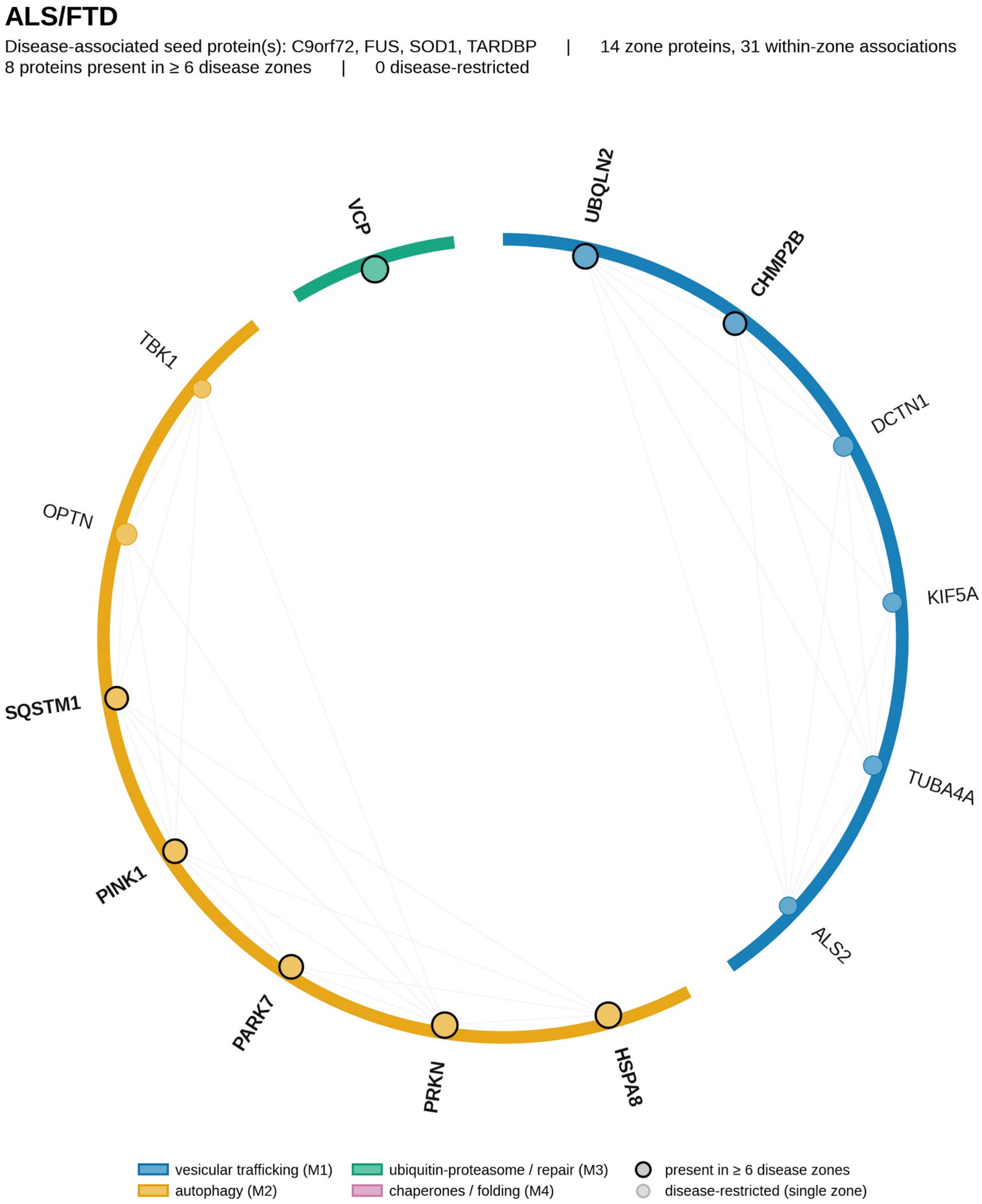

**Figure S8. Complete proteostasis neighborhood of Parkinson’s disease, related to Figures 2 and 3.** Every protein in the primary seed-removed Parkinson’s disease zone (C400; eligible universe; 123 proteins) is shown as a network: nodes are the zone proteins, grouped around the ring by biological process (vesicular trafficking M1, autophagy M2, ubiquitin-proteasome processing and protein repair M3, chaperones and protein folding M4) and colored accordingly, and chords denote within-zone STRING associations (combined score at least 0.4). Proteins present in at least six of the twelve disease zones are ringed and shown in bold; disease-restricted proteins (present in a single zone) are shown in gray. Disease-associated seed protein(s): SNCA. See also Table S2.

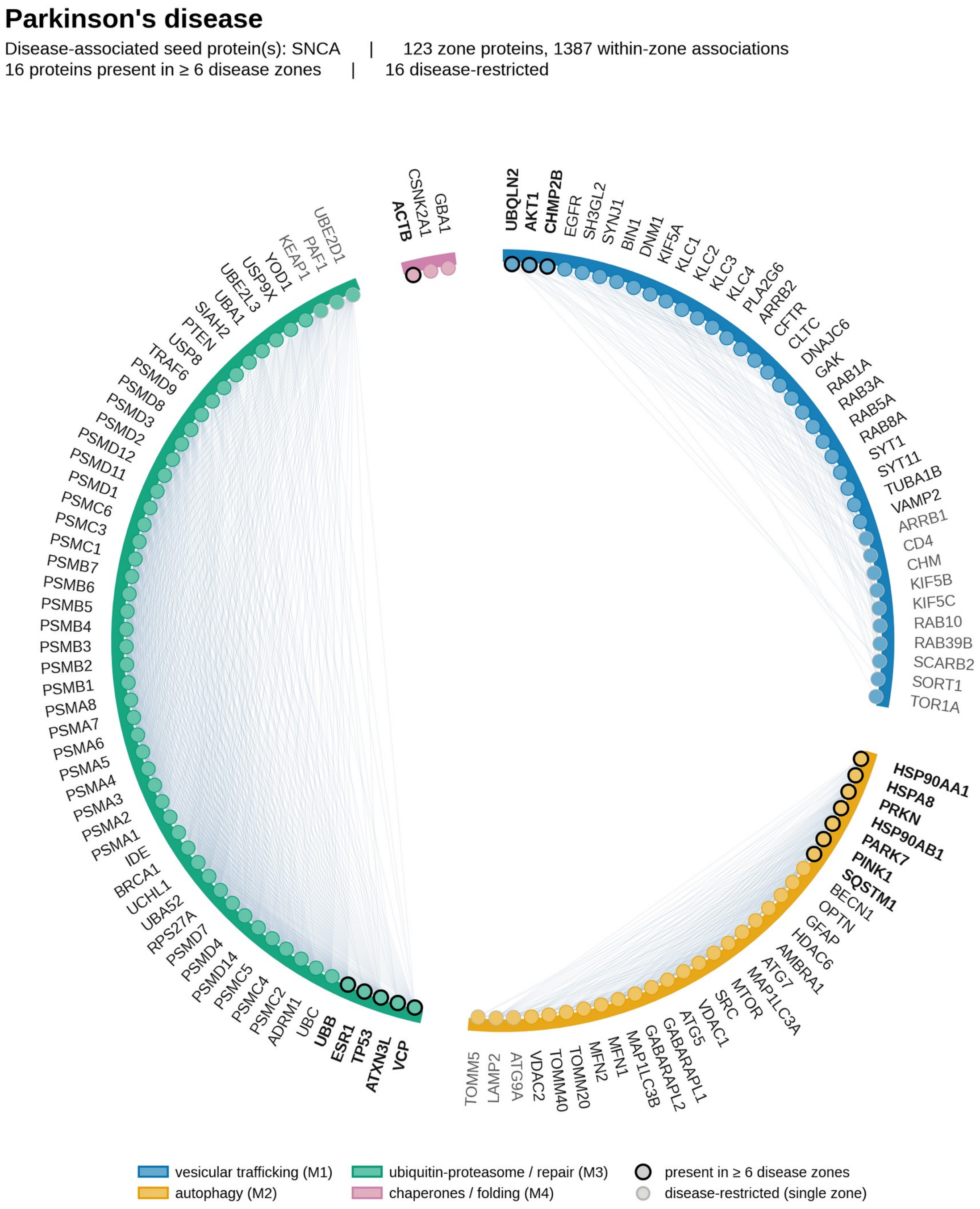

**Figure S9. Complete proteostasis neighborhood of prion disease, related to Figures 2 and 3.** Every protein in the primary seed-removed prion disease zone (C400; eligible universe; 56 proteins) is shown as a network: nodes are the zone proteins, grouped around the ring by biological process (vesicular trafficking M1, autophagy M2, ubiquitin-proteasome processing and protein repair M3, chaperones and protein folding M4) and colored accordingly, and chords denote within-zone STRING associations (combined score at least 0.4). Proteins present in at least six of the twelve disease zones are ringed and shown in bold; disease-restricted proteins (present in a single zone) are shown in gray. Disease-associated seed protein(s): PRNP. See also Table S2.

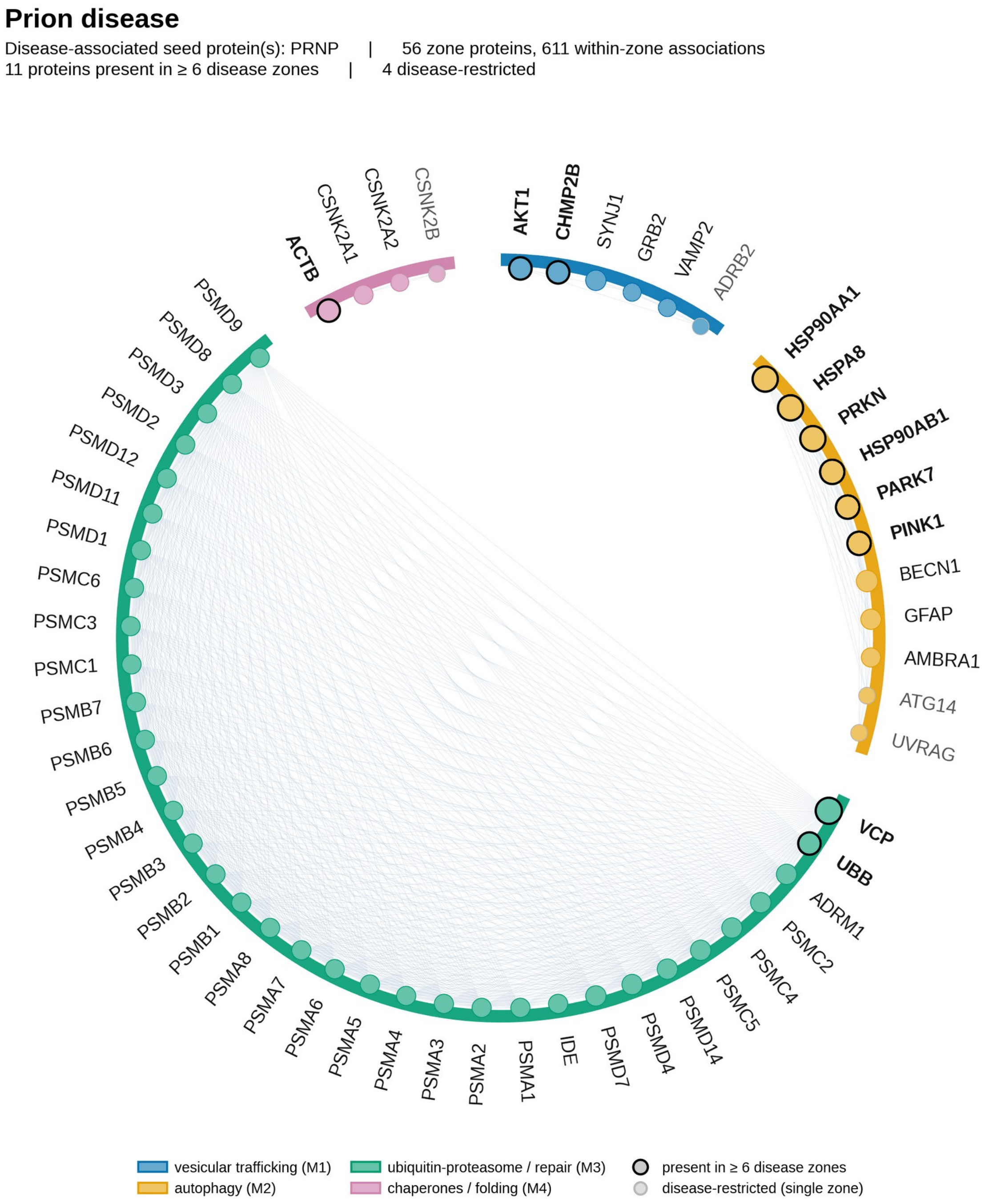

**Figure S10. Complete proteostasis neighborhood of Huntington's disease, related to Figures 2 and 3.** Every protein in the primary seed-removed Huntington's disease zone (C400; eligible universe; 136 proteins) is shown as a network: nodes are the zone proteins, grouped around the ring by biological process (vesicular trafficking M1, autophagy M2, ubiquitin-proteasome processing and protein repair M3, chaperones and protein folding M4) and colored accordingly, and chords denote within-zone STRING associations (combined score at least 0.4). Proteins present in at least six of the twelve disease zones are ringed and shown in bold; disease-restricted proteins (present in a single zone) are shown in gray. Disease-associated seed protein(s): HTT. See also Table S2.

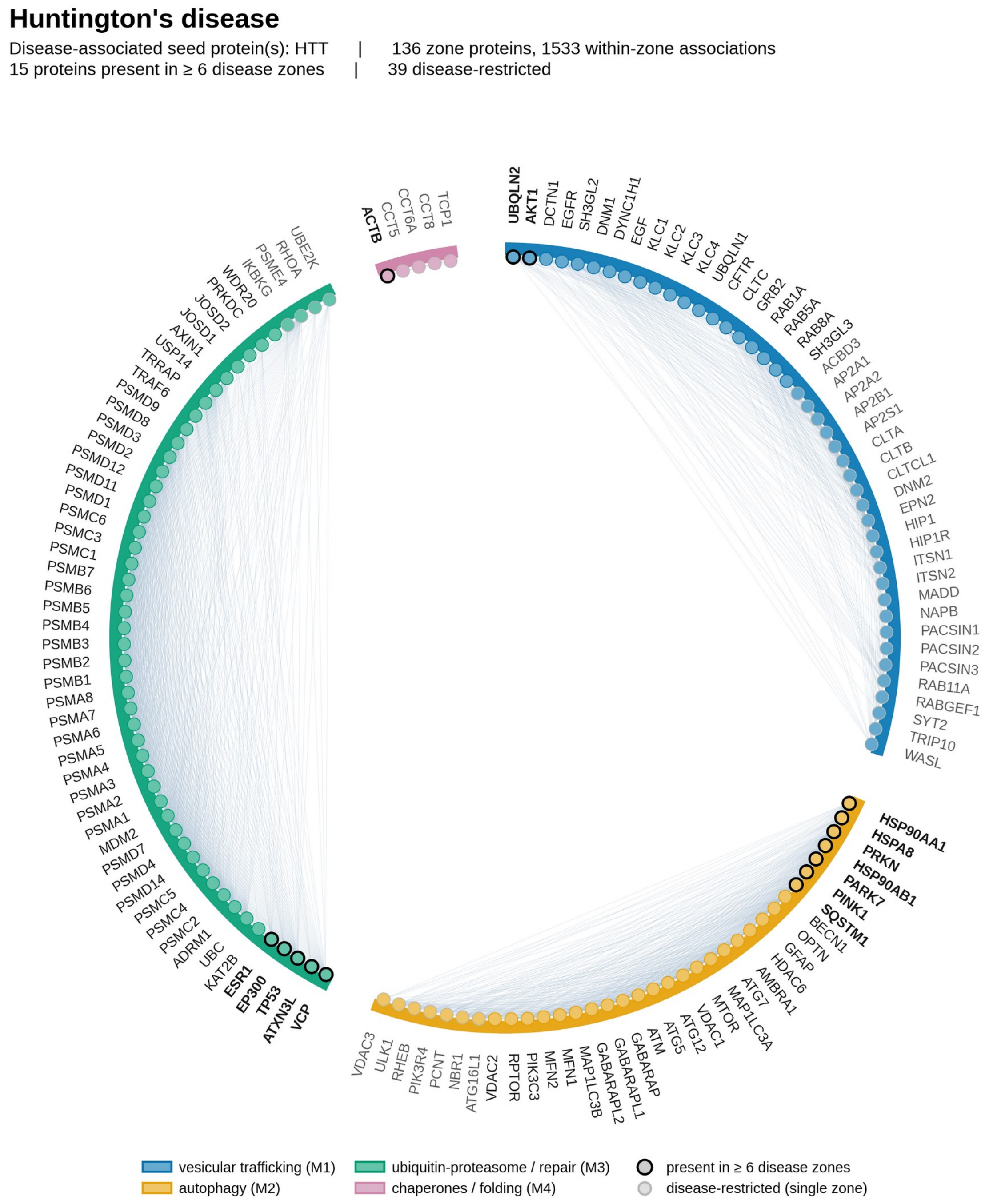

**Figure S11. Complete proteostasis neighborhood of spinal and bulbar muscular atrophy (SBMA), related to Figures 2 and 3.** Every protein in the primary seed-removed spinal and bulbar muscular atrophy disease zone (C400; eligible universe; 96 proteins) is shown as a network: nodes are the zone proteins, grouped around the ring by biological process (vesicular trafficking M1, autophagy M2, ubiquitin-proteasome processing and protein repair M3, chaperones and protein folding M4) and colored accordingly, and chords denote within-zone STRING associations (combined score at least 0.4). Proteins present in at least six of the twelve disease zones are ringed and shown in bold; disease-restricted proteins (present in a single zone) are shown in gray. Disease-associated seed protein(s): AR. See also Table S2.

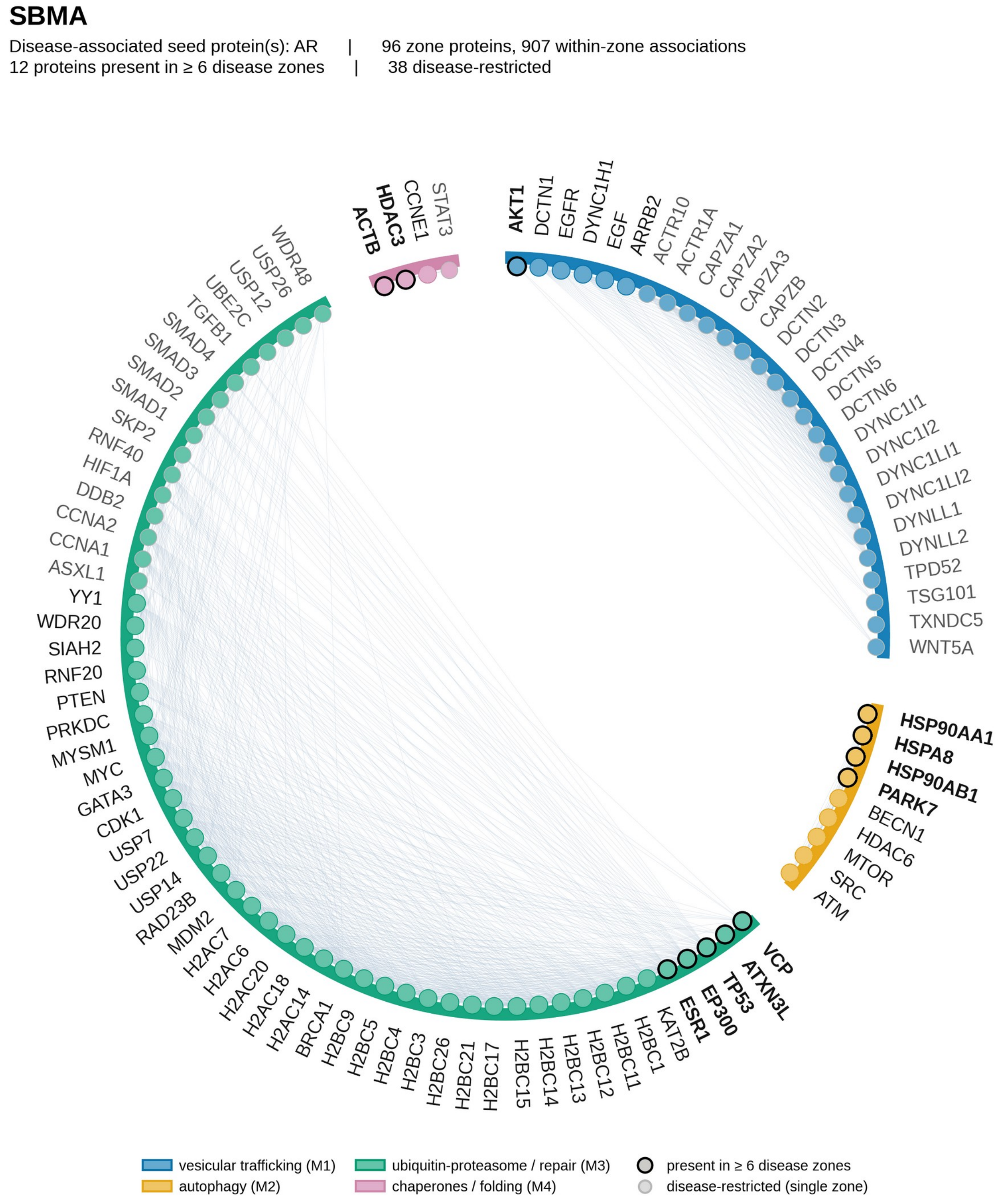

**Figure S12. Complete proteostasis neighborhood of dentatorubral-pallidoluysian atrophy (DRPLA), related to Figures 2 and 3.** Every protein in the primary seed-removed dentatorubral-pallidoluysian atrophy disease zone (C400; eligible universe; 5 proteins) is shown as a network: nodes are the zone proteins, grouped around the ring by biological process (vesicular trafficking M1, autophagy M2, ubiquitin-proteasome processing and protein repair M3, chaperones and protein folding M4) and colored accordingly, and chords denote within-zone STRING associations (combined score at least 0.4). Proteins present in at least six of the twelve disease zones are ringed and shown in bold; disease-restricted proteins (present in a single zone) are shown in gray. Disease-associated seed protein(s): ATN1. This zone is low coverage and is presented for completeness only. See also Table S2.

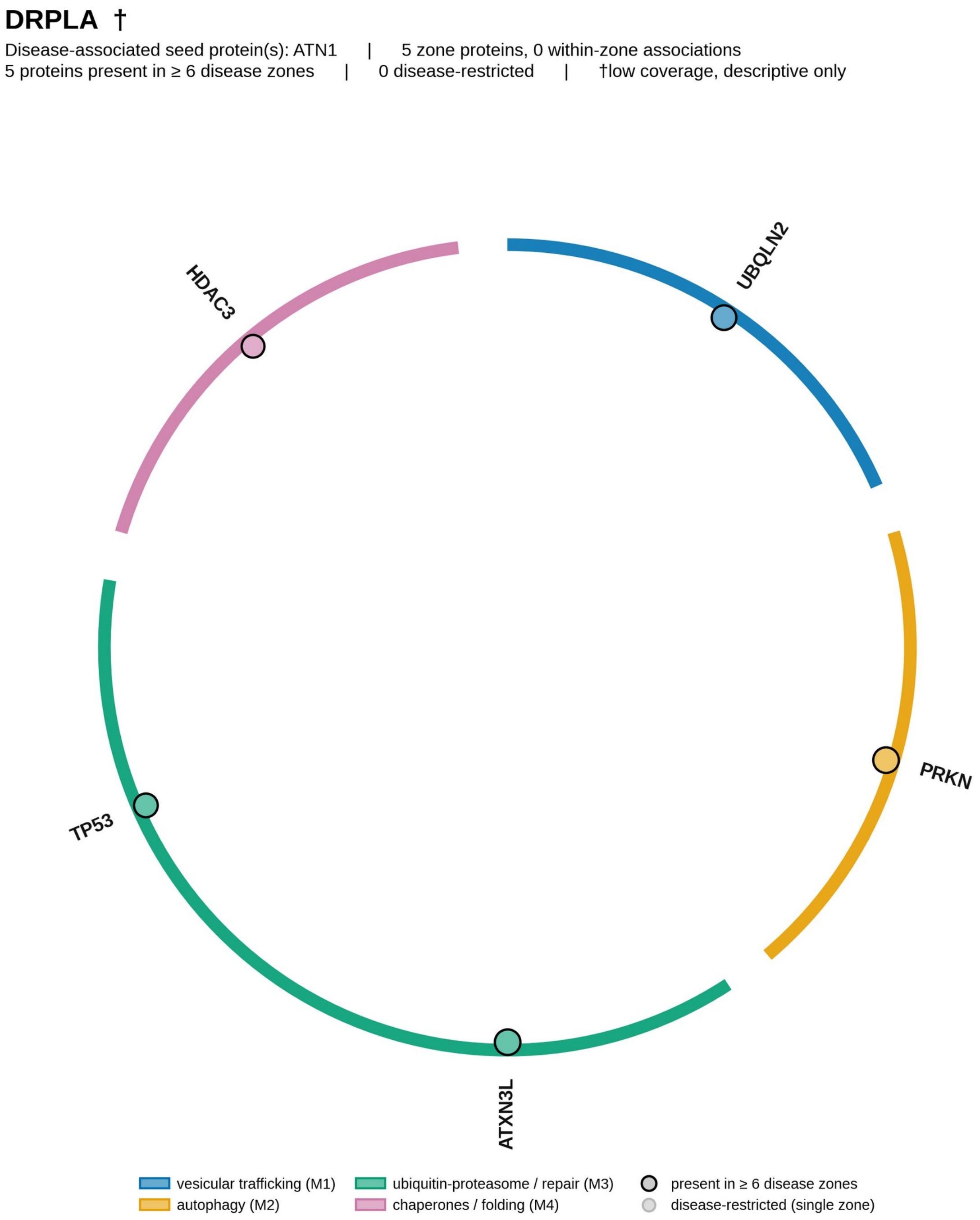

**Figure S13. Complete proteostasis neighborhood of spinocerebellar ataxia type 1 (SCA1), related to Figures 2 and 3.** Every protein in the primary seed-removed spinocerebellar ataxia type 1 disease zone (C400; eligible universe; 14 proteins) is shown as a network: nodes are the zone proteins, grouped around the ring by biological process (vesicular trafficking M1, autophagy M2, ubiquitin-proteasome processing and protein repair M3, chaperones and protein folding M4) and colored accordingly, and chords denote within-zone STRING associations (combined score at least 0.4). Proteins present in at least six of the twelve disease zones are ringed and shown in bold; disease-restricted proteins (present in a single zone) are shown in gray. Disease-associated seed protein(s): ATXN1. See also Table S2.

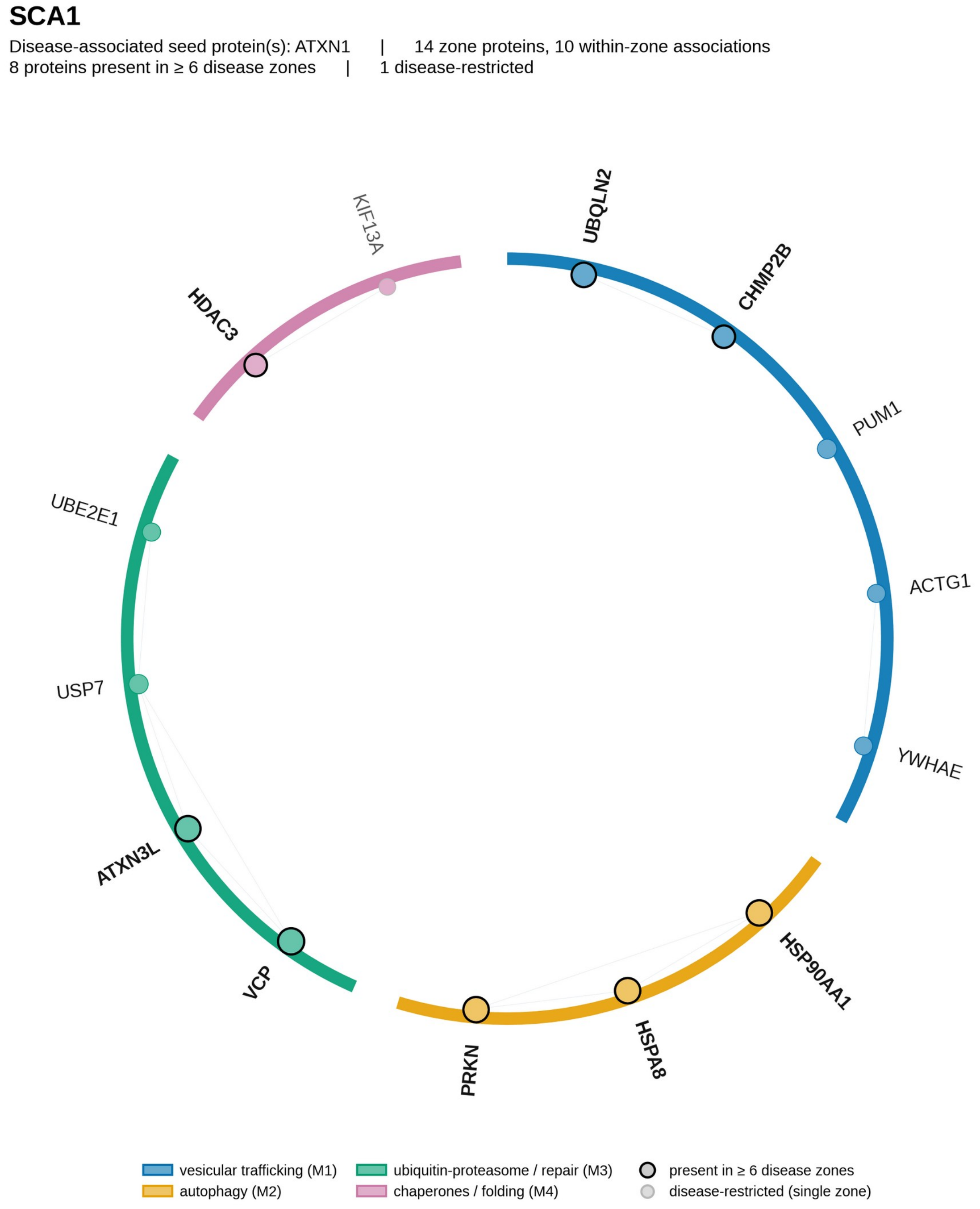

**Figure S14. Complete proteostasis neighborhood of spinocerebellar ataxia type 2 (SCA2), related to Figures 2 and 3.** Every protein in the primary seed-removed spinocerebellar ataxia type 2 disease zone (C400; eligible universe; 33 proteins) is shown as a network: nodes are the zone proteins, grouped around the ring by biological process (vesicular trafficking M1, autophagy M2, ubiquitin-proteasome processing and protein repair M3, chaperones and protein folding M4) and colored accordingly, and chords denote within-zone STRING associations (combined score at least 0.4). Proteins present in at least six of the twelve disease zones are ringed and shown in bold; disease-restricted proteins (present in a single zone) are shown in gray. Disease-associated seed protein(s): ATXN2. See also Table S2.

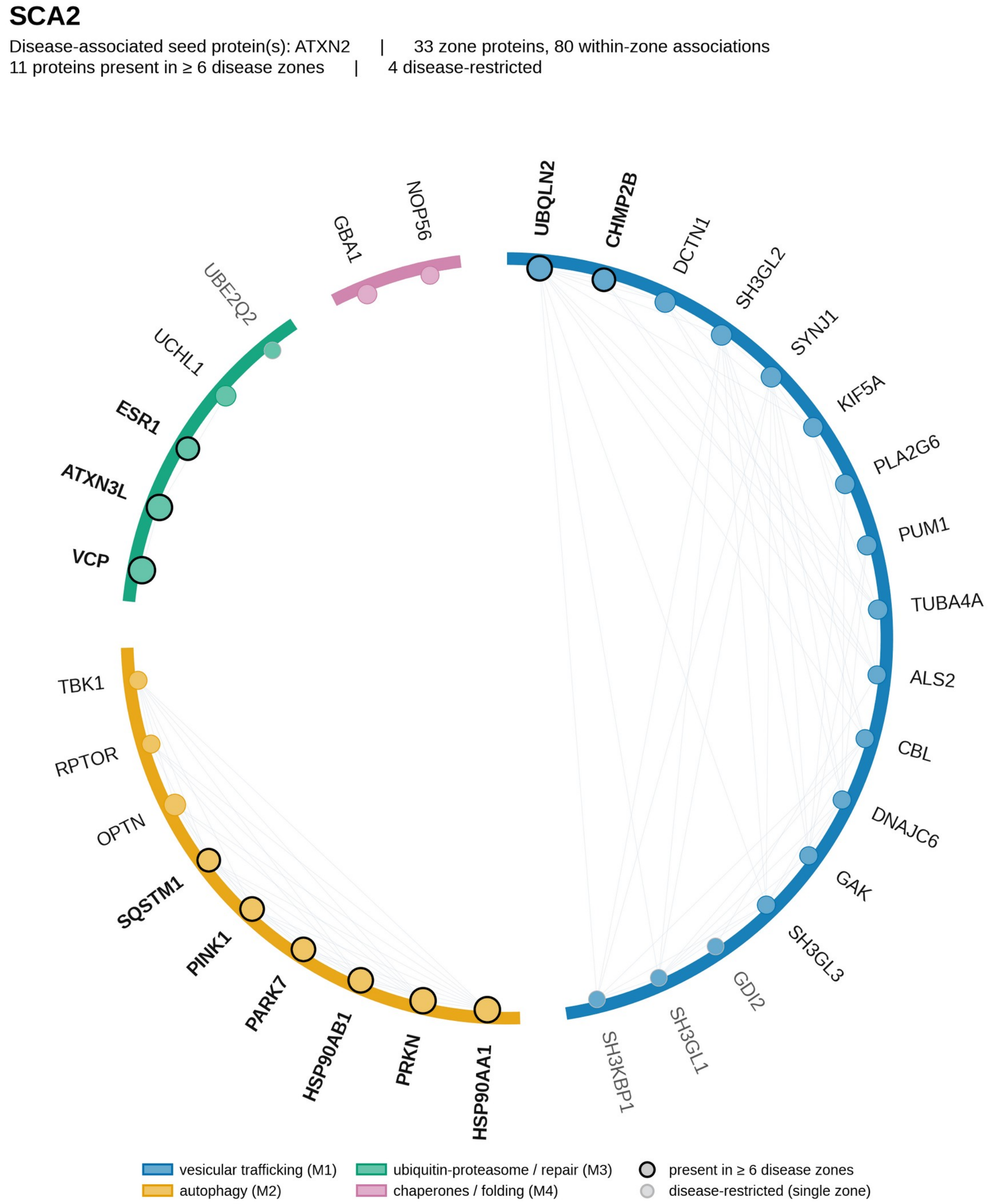

**Figure S15. Complete proteostasis neighborhood of spinocerebellar ataxia type 3 (SCA3, Machado-Joseph disease), related to Figures 2 and 3.** Every protein in the primary seed-removed spinocerebellar ataxia type 3 disease zone (C400; eligible universe; 93 proteins) is shown as a network: nodes are the zone proteins, grouped around the ring by biological process (vesicular trafficking M1, autophagy M2, ubiquitin-proteasome processing and protein repair M3, chaperones and protein folding M4) and colored accordingly, and chords denote within-zone STRING associations (combined score at least 0.4). Proteins present in at least six of the twelve disease zones are ringed and shown in bold; disease-restricted proteins (present in a single zone) are shown in gray. Disease-associated seed protein(s): ATXN3. See also Table S2.

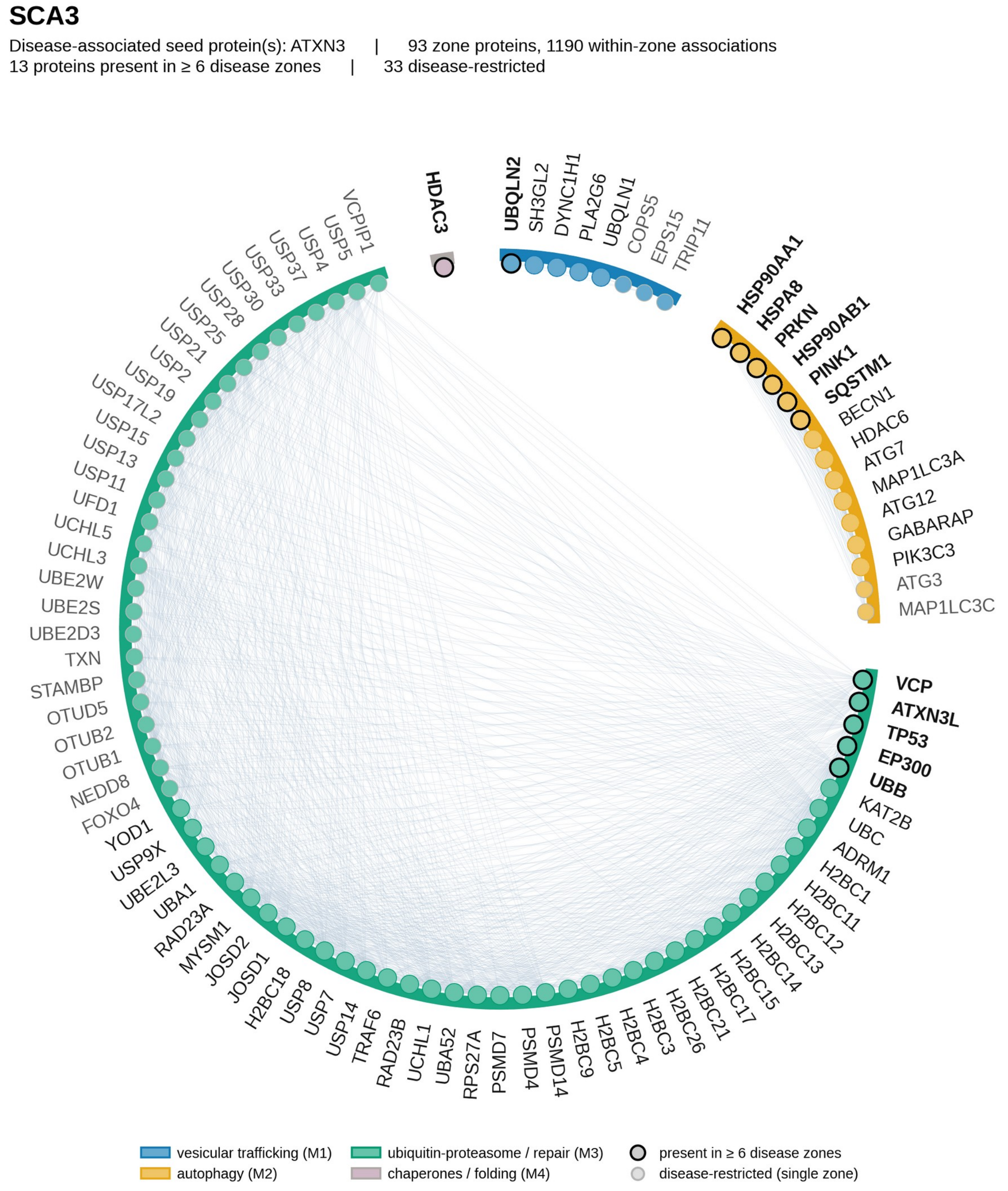

**Figure S16. Complete proteostasis neighborhood of spinocerebellar ataxia type 7 (SCA7), related to Figures 2 and 3.** Every protein in the primary seed-removed spinocerebellar ataxia type 7 disease zone (C400; eligible universe; 47 proteins) is shown as a network: nodes are the zone proteins, grouped around the ring by biological process (vesicular trafficking M1, autophagy M2, ubiquitin-proteasome processing and protein repair M3, chaperones and protein folding M4) and colored accordingly, and chords denote within-zone STRING associations (combined score at least 0.4). Proteins present in at least six of the twelve disease zones are ringed and shown in bold; disease-restricted proteins (present in a single zone) are shown in gray. Disease-associated seed protein(s): ATXN7. See also Table S2.

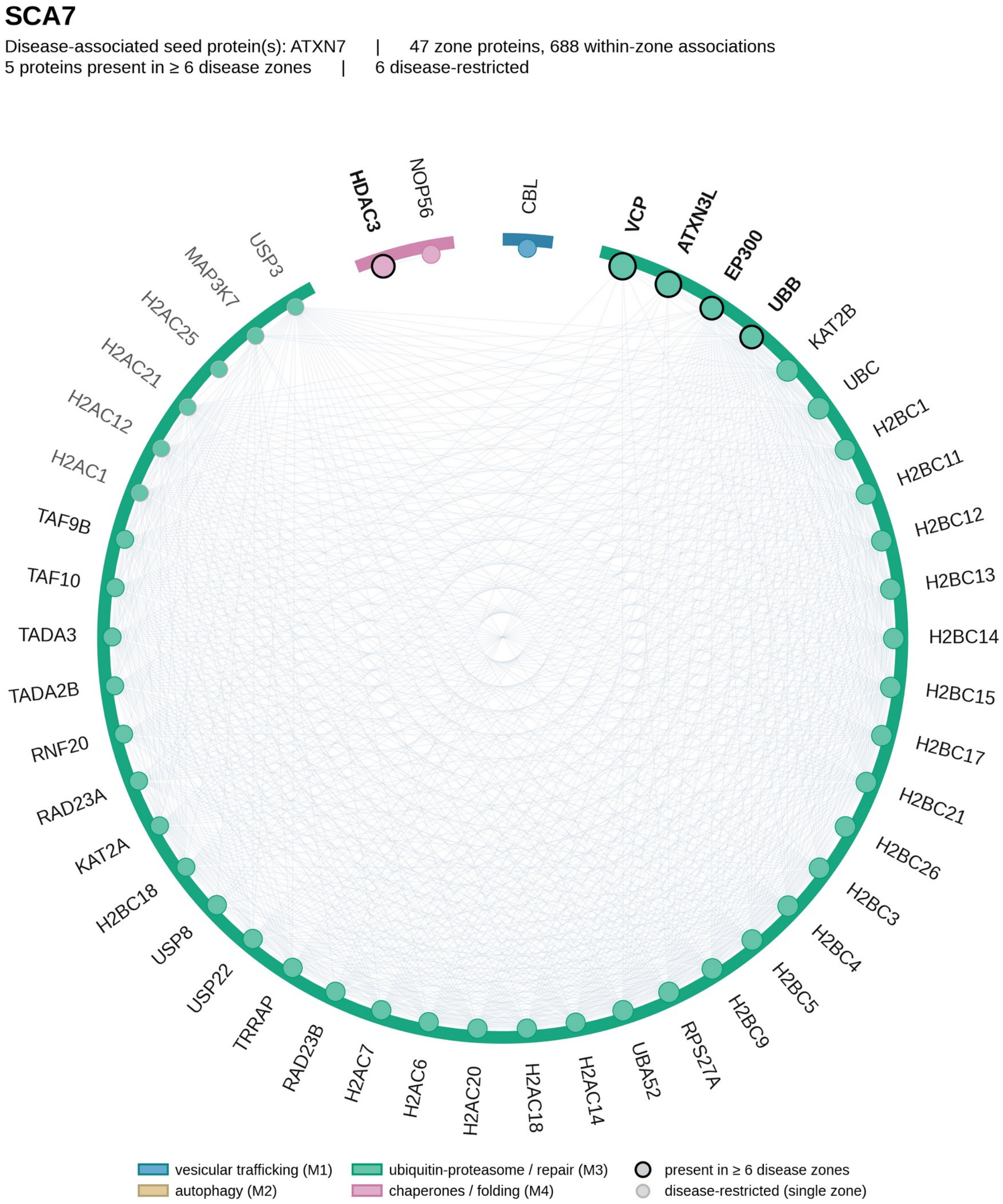

**Figure S17. Complete proteostasis neighborhood of spinocerebellar ataxia type 17 (SCA17), related to Figures 2 and 3.** Every protein in the primary seed-removed spinocerebellar ataxia type 17 disease zone (C400; eligible universe; 74 proteins) is shown as a network: nodes are the zone proteins, grouped around the ring by biological process (vesicular trafficking M1, autophagy M2, ubiquitin-proteasome processing and protein repair M3, chaperones and protein folding M4) and colored accordingly, and chords denote within-zone STRING associations (combined score at least 0.4). Proteins present in at least six of the twelve disease zones are ringed and shown in bold; disease-restricted proteins (present in a single zone) are shown in gray. Disease-associated seed protein(s): TBP. See also Table S2.

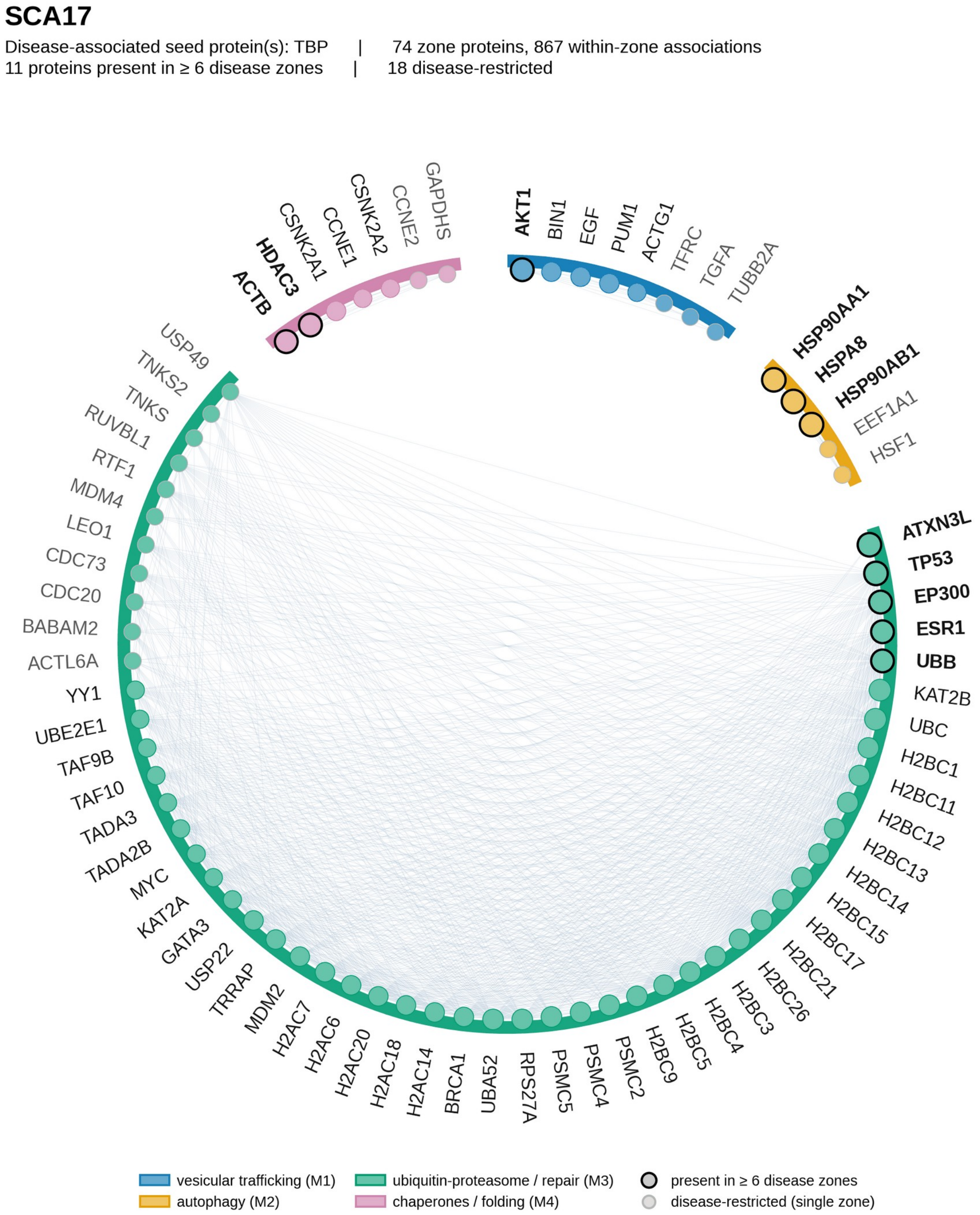
